# HelR is a helicase-like protein that protects RNA polymerase from rifamycin antibiotics

**DOI:** 10.1101/2021.05.10.443488

**Authors:** Matthew D. Surette, Nick Waglechner, Kalinka Koteva, Gerard D. Wright

**Author notes:** Current address: Dalla Lana School of Public Health, University of Toronto (Nick Waglechner).

## Abstract

Rifamycin antibiotics such as rifampin are widely used for the management of tuberculosis and other bacterial infections. These drugs inhibit prokaryotic RNA polymerase (RNAP) by preventing elongation of mRNA resulting in cell death. Rifamycin resistance in the clinic is manifested primarily through amino acid substitutions in RNAP that decrease target affinity for the antibiotics. In contrast, environmental bacteria possess a wide variety of highly specific rifamycin enzyme-mediated resistance mechanisms that modify and inactivate the antibiotics by glycosylation, phosphorylation, ADP-ribosylation, or hydroxylation. Expression of rifamycin resistance is controlled by a common 19bp *cis*-acting rifamycin associated element (RAE) upstream of inactivating genes. Guided by the presence of RAE sequences, we identify an unprecedented ATP-dependent mechanism of rifamycin resistance that acts not by antibiotic inactivation but by protecting the RNAP target. We show that *Streptomyces venezuelae* encodes a helicase-like protein, HelR, which confers broad spectrum rifamycin resistance. Furthermore, HelR is essential for promoting rifamycin tolerance at inhibitory concentrations, enabling bacterial evasion of the toxic properties of these antibiotics. HelR forms a complex with RNAP *in vivo* and rescues transcription inhibition by rifampin *in vitro.* We synthesized a rifamycin photoprobe and demonstrated that HelR directly displaces rifamycins from RNAP. HelR-encoding genes associated with RAEs are broadly distributed in actinobacteria, including many opportunistic *Mycobacterial* pathogens, which cannot currently be treated with rifamycins. This first report of an RNAP protection protein conferring antibiotic resistance and offers guidance for developing new rifamycin antibiotics that can avoid this mechanism.

## Introduction

Rifamycin antibiotics such as rifampin and rifabutin are semisynthetic derivatives of the natural product rifamycin B, discovered in 1957 as a fermentation product of *Amycolatopsis mediterranei* (Sensi, 1983). Rifampin (Rifampicin) emerged in the late 1960s and early 70’s as a frontline treatment for infections caused by mycobacteria, particularly *Mycobacterium tuberculosis* (Floss and Yu, 2005). Rifamycins are potent inhibitors of prokaryotic DNA-dependent RNA polymerase (RNAP) (Wehrli et al., 1968). Bacterial RNAP is comprised of four proteins - α2, β, β′, ω - and a transiently associated σ factor (Korzheva, 2000). Rifamycins bind the β subunit of RNAP (RpoB) and occupy the path where the growing transcript emerges (Campbell et al., 2001). This interaction blocks the RNA exit tunnel impeding the synthesis of mRNA longer than 2-3nt.

A pitfall of rifamycin antibiotics is their high frequency of resistance, approximately 10^-7^ – 10^-9^ per bacterium per cell division (Gillespie, 2002). For this reason, they are primarily used in combination with other antibiotics. Against the slow-growing *M. tuberculosis,* for example, rifampin is frequently combined with isoniazid and ethambutol (WHO, 2008). Genetic studies have identified an 81bp region in *rpoB,* termed the **R**ifampin **R**esistance **D**etermining **R**egion (RRDR), which accounts for ∼95% of all rifamycin resistance mutations in the clinic (Yam et al., 2004). The spectrum of resistance mutations is relatively homogenous, with substitutions in three residues Asp516, His526, and Ser531 (*E. coli* RpoB numbering), which reduce the affinity of rifampin for the RNA exit tunnel and account for ∼85% of rifampin resistant *M. tuberculosis* (Ramaswamy and Musser, 1998).

In contrast to the clinical resistome, the spectrum of rifamycin resistance mechanisms in the environment is highly diverse. While Gram-negative bacteria are primarily intrinsically insensitive due to impermeability and/or active efflux of these compounds, various Gram-positive genera have evolved highly specific mechanisms of rifamycin resistance. Many Actinobacteria, including but not limited to the genera *Streptomyces, Nocardia, Rhodococcus*, and *Mycobacterium,* possess multiple mechanisms of enzymatic inactivation of rifamycins (Dabbs, 1987; Dabbs et al., 1995; Spanogiannopoulos et al., 2012). Four distinct inactivation mechanisms are known, including phosphorylation, ADP-ribosylation, glycosylation, and hydroxylation (**Figure 1)** (Dabbs et al., 1995; Koteva et al., 2018; Spanogiannopoulos et al., 2012, 2014; Surette et al., 2021). Arr enzymes catalyze the transfer of ADP-ribose to the C23 hydroxyl group (**Figure 1A**), thereby sterically blocking this essential hydroxyl that is required for binding to RpoB (Baysarowich et al., 2008). Rgt enzymes use UDP-glucose to glycosylate rifamycins at the same C23 hydroxyl group (**Figure 1A**) (Spanogiannopoulos et al., 2012). Rph enzymes transfer the β- phosphate from ATP to the adjacent C21 hydroxyl group (**Figure 1A**) (Spanogiannopoulos et al., 2014). This hydroxyl group also is essential for productive interaction with RpoB, and the addition of a bulky and negatively charged phosphate group at this position abolishes RNAP binding. On the other hand, the rifamycin monooxygenases (Rox) hydroxylate the naphthoquinone core and inactivate the drug by linearizing the rifamycin macrocycle, thereby destroying the three- dimensional structure of the antibiotic that is required for RNAP inhibition (**Figure 1A)** (Koteva et al., 2018).

**Figure 1.**
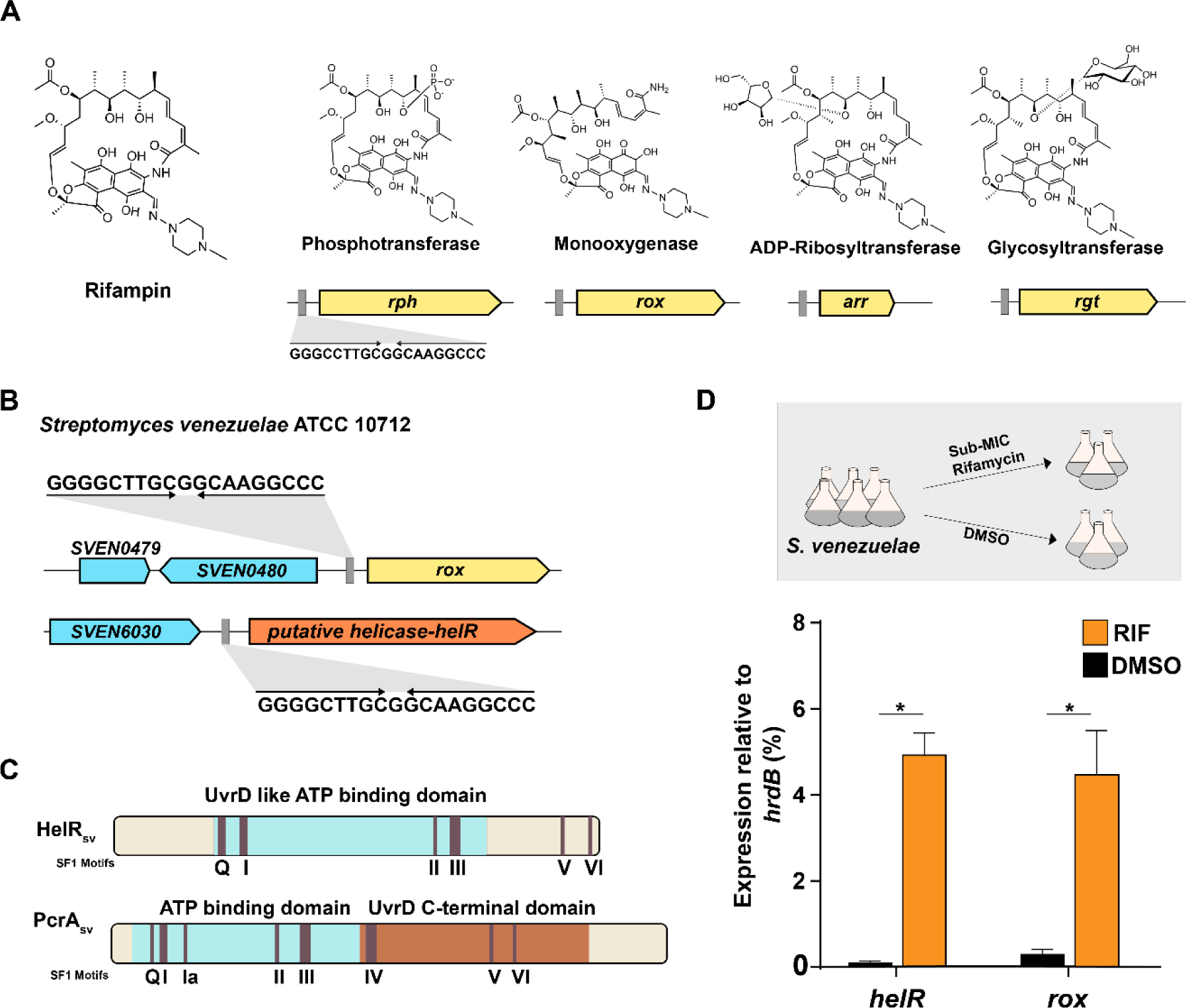
Rifamycin inactivation mechanisms and HelR are controlled by RAE sequences. **A)** Structures of rifampin and its products inactivated by all known group transfer enzymes. The position of the RAE element is shown as a grey rectangle. **B)** RAE sequences are associated with *rox* and *helR* in S. *venezuelae*. **C)** Domain architecture of HelR and model Superfamily 1 helicase PcrA. Conserved Superfamily 1 motifs are depicted. **D)** Expression of *helR* and *rox* normalized to *hrdB* with and without exposure to sub-MIC rifamycins. P values calculated using an unpaired students t-test * P < 0.001.

Groups studying these inactivation mechanisms in the 1990s noted that enzyme production was often inducible by rifamycins (Quan et al., 1997). Over 20 years later, we discovered a 19 bp palindromic sequence upstream from *rgt* (Spanogiannopoulos et al., 2014). This sequence, termed the <u>R</u>ifamycin <u>A</u>ssociated <u>E</u>lement (RAE), was used to identify the gene encoding the enzyme responsible for rifampin phosphorylation in a different *Streptomyces* strain, revealing the RAE’s predictive value in targeting rifamycin resistance genes. Subsequent bioinformatic analysis revealed that RAE sequences are found upstream of all known rifamycin inactivating enzymes (Arr, Rox, Rph, and Rgt) within the Actinobacteria phylum. Furthermore, the RAE was demonstrated to be necessary for the induction of downstream genes in response to rifamycin antibiotics (Spanogiannopoulos et al., 2014). The molecular mechanism underlying this regulation remains unknown; nevertheless, the RAE, and the myriad of inactivating enzymes it controls, are a significant source of rifamycin resistance in Actinobacteria.

While all known rifamycin inactivation mechanisms appear associated with a RAE, the RAE is also found upstream of genes with no known resistance function. A significant proportion of RAEs are upstream of genes annotated as putative helicases (Spanogiannopoulos et al., 2014). Here we demonstrate that the rifamycin associated <u>hel</u>icase-like protein (HelRSv) in *S. venezuelae* is a highly specific rifamycin resistance enzyme. In contrast to all previous RAE-associated genes, HelR is not an antibiotic inactivating enzyme. Instead, HelR directly interacts with RNAP, where it displaces bound rifamycins, thereby relieving inhibition. We also examine the distribution of HelR and the closely related HelD proteins, which are abundant in the genomes of Firmicutes and Actinobacteria. HelR homologs associated with RAEs fall into distinct protein clusters, indicating that only a small subset of these proteins are likely involved in rifamycin resistance.

## Results

### *helR* is a rifamycin inducible resistance gene in *S. venezuelae*

In addition to its ubiquitous presence upstream of genes encoding rifamycin inactivating enzymes, the RAE is also associated with genes of unknown function. *Streptomyces venezuelae* ATCC10712 is a model *Streptomyces* that has two RAEs in its genome (**Figure 1B**). One is associated with a rifamycin monooxygenase (Rox), and the other with a putative helicase (*helR*). HelRSv is homologous to superfamily 1 helicases (**Figure 1C**), a protein class not previously associated with antibiotic resistance. Helicases are ssDNA translocases that couple ATP hydrolysis with directional movement along the DNA strand, where they collide with dsDNA and separate individual strands. While helicases are essential for DNA replication, they are not required for transcription (Dangkulwanich et al., 2014; Dillingham and Dillingham, 2011; Singleton et al., 2007). RNAP can melt promoter DNA to form a transcription bubble and open downstream DNA as the bubble moves during transcription without a helicase. Consequently, it was not apparent how the unwinding of any specific DNA or RNA segment could impact rifamycin activity, warranting genetic control by a RAE.

HelR is most similar to superfamily 1 helicases defined by core structural similarities and characteristic amino acid motifs (Fairman-Williams et al., 2010). We were able to identify many of these motifs in HelRSv (**Figure 1C**). However, when we compared the domain architecture of HelRSv to well-studied SF1 helicases like UvrD/PcrA using Interpro (Mitchell et al., 2019), we noticed that HelR lacks 1 of the two core SF1 helicase domains (**Figure 1C**). HelR is predicted to have a UvrD-like ATP binding domain that makes up the middle of the protein but lacks a UvrD- like C-terminal domain required for DNA binding. Without this domain, it is unclear how HelR could function as a helicase, and we hypothesized that it has a different function associated with rifamycin antibiotic activity.

We first confirmed the expected rifamycin-dependent expression of *helR* and *rox* using RT-qPCR (**Figure 1D**). RNA was isolated after a two-hour incubation with 0.5 µg/mL rifamycin SV (1/16X Minimal Inhibitory Concentration (MIC)) or DMSO (as vehicle control). The levels of *helR* and *rox* transcript were normalized to the constitutively expressed housekeeping gene *hrdB.* Both *helR* and *rox* showed low-level expression in the absence of rifamycin and were respectively induced 50-fold and 15-fold (P<0.001) in the presence of the antibiotic. Higher basal expression of *rox* appears to be the source of the difference in magnitude of induction, as both reach similar levels of maximal mRNA expression. Consistent with the presence of the RAE in its promoter region, *helR* is induced by rifamycins in *S. venezuelae*.

We generated single gene deletions of Δ*rox* and Δ*helR* as well as a double deletion of Δ*helR*Δ*rox* in *S. venezuelae* to explore the individual effects of both RAE-associated genes. HelRSv conferred robust resistance to all rifamycins tested (rifamycin SV, rifampin, rifabutin, and rifaximin), with increases in susceptibility ranging from 8- to 16-fold upon deletion (comparing Δ*rox* to Δ*helR*Δ*rox*) (**Table 1**). *S. venezuelae* Δ*rox* was 4-8-fold more sensitive, except against rifabutin, an exceedingly poor substrate for Rox (Koteva et al., 2018). Surprised that *helR* confers higher resistance levels than *rox*, we confirmed that *helR* is not required for induction of *rox* (**Figure S1**). The MIC of rifampin for *S. venezuelae* falls >30-fold from an already low value of 0.5 µg/ml to 0.016 µg/mL in *S. venezuelae* Δ*helR*Δ*rox,* highlighting the efficacy of these dual resistance mechanisms. None of the strains tested showed altered susceptibility to fidaxomicin, an antibiotic that also targets RNAP but with a different binding site and mechanism of action than rifamycins (Lin et al., 2018). Vancomycin and tetracycline, which respectively target cell wall biosynthesis and translation (Van Bambeke et al., 2004; Dürckheimer, 1975), also showed no difference in susceptibility. We could complement the *helR* mutant by cloning the gene into pIJ10257, an integrative vector that drives expression using the high-level constitutive promoter P*ermE\** (Hong et al., 2005). For all rifamycins, *S. venezuelae* Δ*helR*Δ*rox* pIJ:*helR* returned to the MIC levels of *S. venezuelae* Δ*rox* and did not rise above it, indicating that overexpression of *helR* does not confer additional resistance.

**Table 1.**
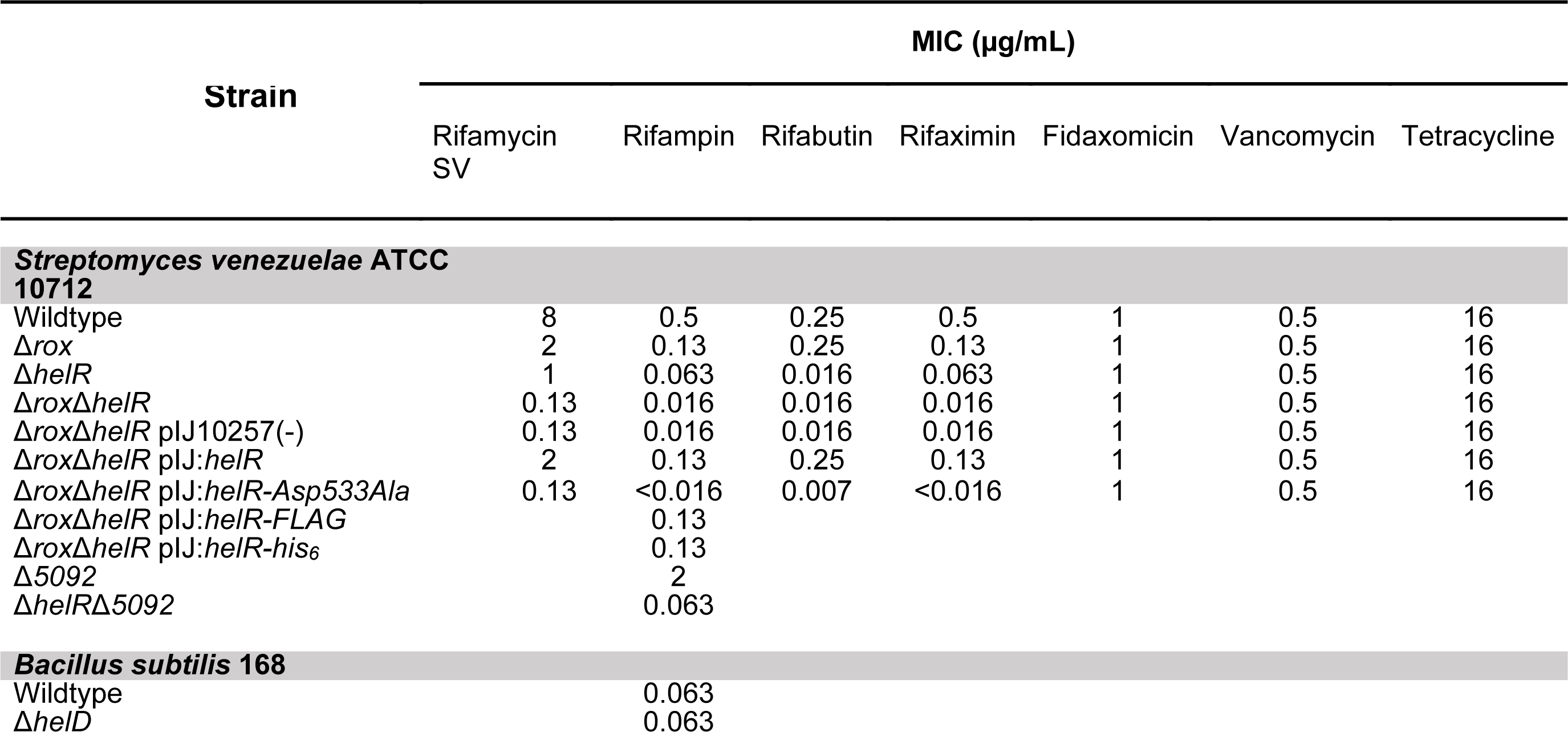
Susceptibility testing

The presence of conserved SF1 helicase motifs required for ATP binding and hydrolysis in HelR suggests that it has ATPase activity, which might be necessary for antibiotic resistance. Helicase motifs I and II (**Figure 1B**) correspond to the Walker A and B motifs found in many proteins that bind and hydrolyze ATP (Fairman-Williams et al., 2010). A conserved Asp and Glu in motif II are responsible for coordinating an essential catalytic Mg^2+,^ and the substitution of either of these amino acids abolishes ATPase activity in other helicases (Raney et al., 2013). We prepared an ATPase-impaired mutant by substituting the Asp of HelR’s motif II (Val532<u>AspGlu</u>AlaGln) to Ala. This mutant did not rescue resistance in *S. venezuelae* Δ*rox*Δ*helR* against all rifamycins tested (**Table 1**). In the case of rifampin, rifabutin, and rifaximin, HelRSvAsp533Ala made cells slightly more susceptible to the antibiotic. The ATPase activity of HelR is therefore required for rifamycin resistance.

Having established that HelR is a novel ATP-dependent rifamycin resistance enzyme, we turned to elucidating its function. All previously characterized genes found under the control of a RAE encode rifamycin inactivating enzymes, so we investigated this possibility for HelR. To monitor the inactivation of rifamycins, mycelia from 24-hour old cultures of *S. venezuelae* were washed and resuspended in fresh media containing 5 µg/ml of rifamycin SV and incubated for 24 hours to allow sufficient time for antibiotic inactivation. Culture supernatant was then applied to a lawn of rifamycin sensitive *Bacillus subtilis* 168 to assess whether the antibiotic had been inactivated (**Figure 2A**). Antibiotic activity was lost from wild-type *S. venezuelae* and Δ*helR* strains, consistent with inactivation of rifamycin SV due to expression of *rox.* On the other hand, *S. venezuelae* Δ*rox* and Δ*helR*Δ*rox* produced zones comparable to the media control (rifamycin SV without any *S. venezuelae*), indicating that Rox is the sole inactivating enzyme in *S. venezuelae* and that HelR must confer resistance through another mechanism.

**Figure 2.**
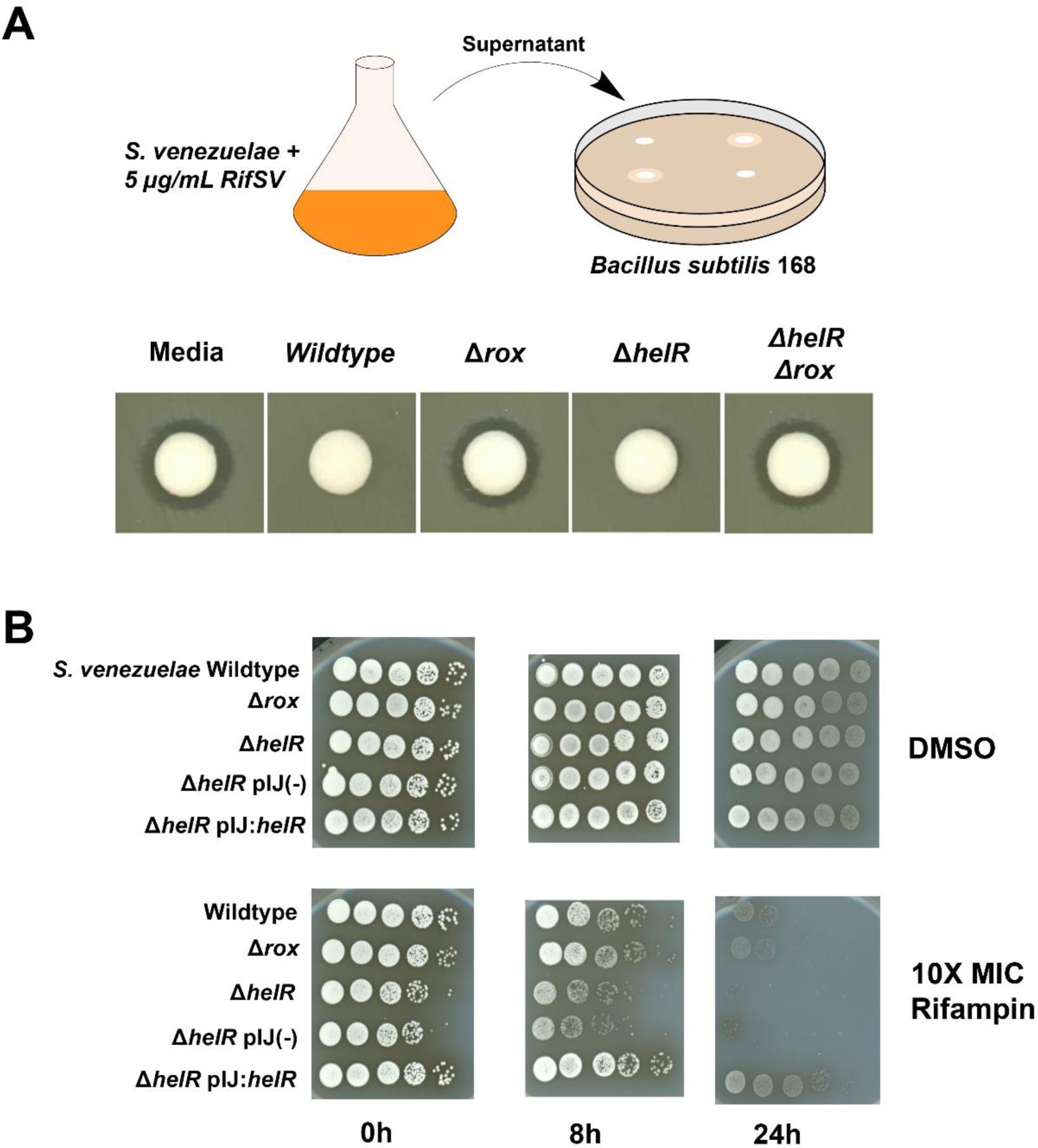
A HelR is not an inactivating enzyme but confers tolerance to rifamycins. **A)** *S. venezuelae* cultures were incubated for 24 hours in TSB + 5µg/mL rifamycin SV. The supernatant of this culture was applied to a cellulose disc and placed on a lawn of indicator bacteria (*B. subtilis*) to monitor antibiotic inactivation. Media control contained only TSB and rifamycin SV. **B)** *S. venezuelae* in exponential growth phase were standardized by OD600nm and exposed to 10X the strains respective MIC of rifampin. Cells were removed by centrifugation and washed several times to remove all remaining rifampin at the noted timepoints. Serial 10- fold dilutions were spotted onto Bennett’s agar to assess viability (dilutions are plated left to right).

### HelR confers tolerance to rifamycins

When conducting disc diffusion assays with rifamycins and *S. venezuelae* mutants, we observed that strains lacking *helR* grew with very sharp margins around their zones of inhibition. In contrast, the wildtype and Δ*rox* strains had gradients of decreasing growth towards the disc. If these plates were incubated for long periods, *helR^+^* strains would eventually grow within the initial zone of inhibition while *helR^-^* strains would not. We interpreted this phenotype as antibiotic tolerance and studied it further. Rifamycins are bactericidal towards Gram-positive bacteria and especially mycobacteria (Floss and Yu, 2005). We performed time-kill experiments on exponentially growing *S. venezuelae* with rifampin concentrations 10X the MIC of each strain. Cells were recovered and resuspended in fresh media three times to remove the remaining antibiotic, and tenfold dilutions were spotted onto agar to assess viability (**Figure 2B**). Even at the 0 h timepoint where cells were only momentarily exposed to rifampin, *S. venezuelae* Δ*helR* almost immediately lost ∼10-fold viability. In contrast, wildtype *S. venezuelae*, Δ*rox,* and Δ*helR* pIJ:*helR* were unaffected. This difference became more pronounced after 8 hours of antibiotic exposure, with Δ*helR* losing closer to 100-fold viability. Wildtype and Δ*rox* show a very slight decline in viability at this timepoint, whereas Δ*helR* pIJ:*helR* was fully viable. At 24 hours, a significant loss of viability for wildtype and Δ*rox* cells is observed, although they are still orders of magnitude more viable than *helR.* At this timepoint, Δ*helR* pIJ*helR* is significantly more viable than wildtype cells. While overexpression of *helR* does not lead to a rise in MIC, it does increase drug tolerance. In addition to being an effective rifamycin resistance enzyme, these data show that HelR also allows *S. venezuelae* to tolerate inhibitory concentrations of these drugs for longer periods of time.

### HelR forms a complex with RNA polymerase

*Bacillus subtilis* HelD has the same overall domain architecture as HelRSv consisting of a core UvrD-like ATP binding domain but lacking a C-terminal DNA binding domain, yet HelRSv shares only minimal amino acid conservation with HelD (18% identity, 32% similarity). HelD lacks helicase activity *in vitro,* but it does bind RNAP (Wiedermannovà et al., 2014). Furthermore, HelD stimulates transcription *in vitro* in an ATP-dependent manner (Wiedermannovà et al., 2014). UvrD, a well-characterized and broadly conserved SF1 helicase, has also been reported to interact directly with RNAP during transcription-coupled repair (Epshtein et al., 2015). Based on this precedent, we hypothesized that HelR binds RNAP and that this interaction is linked to rifamycin resistance. We purified native RNAP from *S. venezuelae* grown in the presence of sub-MIC rifamycin (RIF+) and without (RIF-). We used the identical growth conditions used for the RT-qPCR experiments (**Figure 1D**) to ensure induction of *helR*, and we quantified the proteins in each sample by LC MS-MS (**Figure 3A**). A total of 76 proteins were identified with high confidence, most of which did not change significantly in abundance between samples. HelRSv, in contrast, was the most enriched protein (426-fold) following rifampin exposure (**Figure 3A, Supplemental File 1, Table S1**). Indeed, when purified RNAP fractions were analyzed by SDS-PAGE, a prominent band was visible at ∼80kDa consistent with the size of HelRSv(**Figure 3B**). This band was excised from the gel and confirmed to be HelRSv using LC MS-MS. We did not detect Rox in our samples, suggesting that the association of HelRSv with RNAP is not an artifact of overexpression since this protein is also induced by rifamycin exposure.

**Figure 3.**
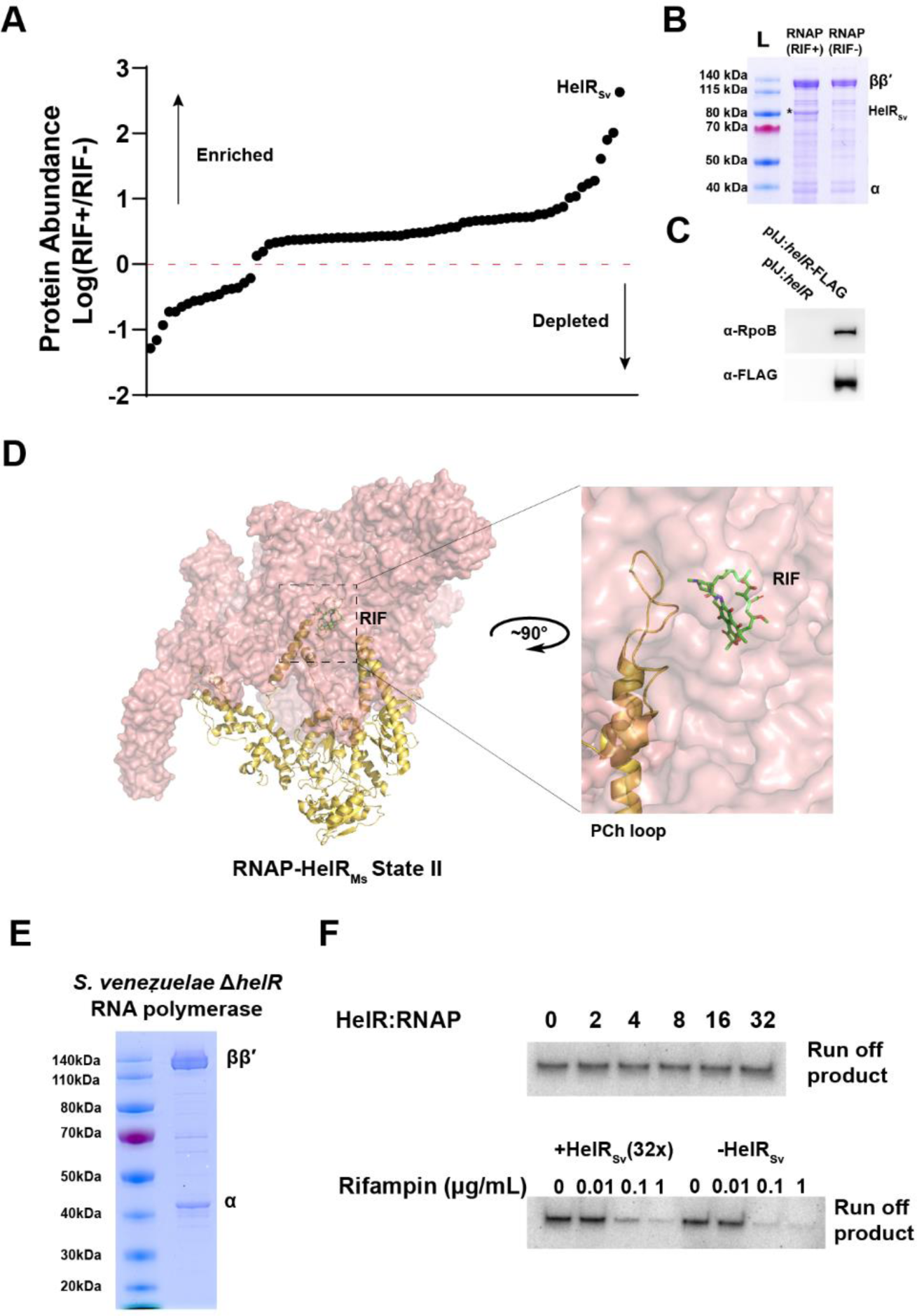
HelRSv co-purifies with RNA polymerase in *S. venezuelae* and blocks rifampin activity *in vitro*. **A)** Quantitative proteomic comparison of RNA polymerase purifications from rifamycin induced (RIF+) or non-induced (RIF-). Raw abundance data are expressed as the log of the ratio of RIF+ divided by RIF- and plotted in rank order. **B)** SDS-PAGE of RNAP preparations from induced and non-induced cells. The RNAP β, β’ and α subunits are labelled. *denotes the band corresponding to HelRSv **C)** Immunoprecipitation of soluble protein from *S. venezuelae* constitutively expressing native HelRSv or HelRSv-FLAG using α-FLAG resin. Protein was eluted from the resin using FLAG peptide and probed for the presence of RNAP (using an antibody which was raised against the β-subunit) and HelR (α-FLAG). **D)** Alignment of HelRMs:RNAP complexes from *M. smegmatis* (PDB IDs 6XYU and 6YYS for states I and II respectively) with a rifampin bound structure of *M. smegmatis* RNAP (6CCV) was used to model rifampin into the HelRMs:RNAP complexes. **E)** SDS-PAGE of native RNA polymerase isolated from *S. venezuelae* Δ*helR* β, β’, and α subunits are labelled. **F) (top)** Autoradiograph of transcripts from multiple round *in-vitro* transcription reaction with increasing molar ratio of HelRSv:RNAP(σ^hrdB^) **(bottom)** Multiple round transcription reactions performed in the presence of rifampin (RIF) in the presence and absence of a large molar excess of HelRSv.

Since purification of RNAP yields HelRSv, we reasoned that the reciprocal experiment, purification of HelRSv from *S. venezuelae,* should yield RNAP. We constructed a C-terminal FLAG-tagged HelRSv, expressed this protein in *S. venezuelae* Δ*helR,* and found that this construct restored wild-type levels of rifampin resistance and was therefore functional (**Table 1**). We performed co-immunoprecipitation using α-FLAG resin from soluble proteomes of *S. venezuelae* Δ*helR* constitutively expressing either native HelRSv or HelRSv-FLAG (**Figure 3C**). Western blotting confirmed the presence of HelRSv and RNAP from cells expressing HelRSv-FLAG and an absence of RNAP in cells expressing tag-free HelRSv. These data show that RNAP and HelRSv form a stable complex *in vivo*. Because HelRSv-FLAG was expressed constitutively and co-precipitated with RNAP, the presence of rifamycin is not required for complex formation. The association of HelRSv with RNAP in wildtype *S. venezuelae* is the consequence of rifamycin- mediated HelRSv induction. The formation of a complex with HelRSv and the molecular target of rifamycin antibiotics lead us to hypothesize that HelR may function as a protection protein that either prevents or reverses rifamycin binding in an ATP-dependent fashion.

While writing this manuscript, we became aware of three co-structures of HelD in complex with RNAP published simultaneously; two describing HelD from *Bacillus subtilis* (HelDBs) and the third from *Mycobacterium smegmatis*, which has 35% amino acid identity with HelRSv (Kouba et al., 2020; Newing et al., 2020; Pei et al., 2020). The *M. smegmatis* protein is referred to as HelDMs and is purported to be functionally equivalent to HelDBs. However, we observed that *helDMs* is associated with a RAE. Furthermore, this protein is among the most abundant in the cell following exposure to sub-MIC rifampin, and deletion of this gene results in increased susceptibility to rifampin (Hurst-Hess et al., 2019). We predict that this protein is functionally equivalent to HelRSv and should be renamed HelRMs to avoid confusion with genuine HelDs, which have no known role in inducible rifamycin resistance. Both HelDBs and HelRMs bind equivalent sites on RNAP, and both RNAP complexes appear incompatible with DNA in the primary channel. They both “wedge” open the β’ Clamp weakening the interaction between RNAP and DNA and project appendages deep into RNAP. Additionally, both proteins possess a secondary channel arm (SCA) which are structurally similar to transcription factors such as GreA/B (Kouba et al., 2020; Pei et al., 2020). The SCA in HelDBs extends into and occupies the active site. On the other hand, HelRMs has a shorter SCA, which does not enter the active site; instead, this protein possesses a primary channel loop (PCh). Three distinct states of the HelRMs-RNAP complex were solved by Kouba *et al*.. The conformation of the PCh loop was found to be disordered in state I, but in state II, it had folded into the primary channel and interacts with the catalytic Mg^2+^ and mobile domains in the active site. We noted that the path of the PCh loop comes very close to the rifamycin binding pocket on RpoB (**Figure 3D**). The PCh loop does not directly contact the rifamycin binding pocket, nor does this pocket appear to be distorted in the structure of State II. Regardless, the existence of this appendage on a HelD-like enzyme that specifically confers rifamycin resistance is unlikely to be coincidental. Consistent with our hypothesis that HelRSv is an RNAP protection protein, HelRMs possesses an appendage uniquely suited for dislodging rifamycins from their target.

### Mechanism of action of HelR

To test our hypothesis that HelR is a rifamycin protection protein, we reconstituted transcription *in vitro* using RNAP purified from *S. venezuelae* Δ*helR* to ensure that trace amounts of HelRSv did not contaminate preparations. To complete the reagent requirements for *in vitro* transcription, we purified the housekeeping sigma factor for *Streptomyces* σ^HrdB^ and synthesized a modified PermE* with only one transcription start site (Tss) (**Figure S2**). All attempts to express recombinant HelRSv with various affinity and solubility tags in *E. coli* were unsuccessful. To overcome this problem, we constructed HelRSv with a C-terminal His6 tag and expressed this protein in *S. venezuelae*. As with the C-terminal FLAG tag, this construct completely restored rifampin resistance, indicating normal function (**Table 2**). We were able to purify His6-tagged HelRSv directly from *S. venezuelae* for use in *in-vitro* assays.

Examination of HelDBs and HelRMs :RNAP complexes suggest that binding of DNA and HelD are mutually exclusive, which means that RNAP can be engaged with DNA or HelD, but not both. Somewhat paradoxically, HelDBs can stimulate transcription in multiple round assays by removing stalled transcription complexes from template DNA. Interestingly HelRMs has been shown to possess the ability to remove stalled elongating complexes, but there is no reported evidence that this protein can stimulate transcription *in vitro* (Kouba et al., 2020). We first added HelRSv in increasing concentrations to determine if this protein had any general effects on *in vitro* transcription (**Figure 3E**). In multi-round assays, we found that HelRSv had no discernable impact on the amount of transcript formed, even at a 32-fold molar excess over RNAP. The inability of HelRSv to stimulate transcription in this context is further evidence that HelR is functionally distinct from HelD*Bs* and is instead a dedicated resistance enzyme. In agreement with our hypothesis that HelR functions to protect RNAP from rifamycins, we show that the addition of HelRSv to *in vitro* transcription reactions offers protection from inhibitory concentrations of rifampin (**Figure 3F**).

To better understand the molecular mechanism of HelR-mediated rifamycin resistance, we sought a more direct method to measure the presence of rifamycins bound to RNAP. Consequently, we designed and synthesized a rifamycin photoaffinity-probe (RPP) (**Figure 4A, Figure S3**). We used rifamycin B as a scaffold to take advantage of its free carboxylic acid in a location on the antibiotic known to be tolerant to substitutions. We coupled a benzophenone-alkyne to rifamycin B through the carboxylic acid using standard methods (**Figure S3**). Excitation of the benzophenone by long wavelength UV light (365 nm) generates a highly reactive carbene that can crosslink to nearby proteins (Murale et al., 2017) . Lastly, we used ‘click chemistry’ to link the rifamycin-benzophenone to commercially available biotin-PEG3-azide, allowing us to detect the presence of the crosslinked probe using a streptavidin-HRP conjugate (Kolb et al., 2001). RPP recapitulates all relevant properties of a rifamycin antibiotic. It inhibits the growth of both *B. subtilis* 168 and *Streptomyces venezuelae*; a known rifamycin resistance substitution in RpoB (*B. subtilis* numbering, H482Y) confers RPP resistance; and importantly, *helR* also provides increased resistance to RPP (**Figure 4B**). We used RPP to photolabel purified *S. venezuelae* RNAP *in vitro.* Samples were exposed to increasing concentrations of RPP, proteins were separated by SDS-PAGE, transferred to PVDF membranes, and probed with streptavidin-HRP (**Figure 4C**). Blotting of purified RNAP revealed two endogenously biotinylated proteins present in our preparations; this had no impact on our ability to label and quantify labeling of RNAP with RPP (**Figure S4**). We observed concentration-dependent labeling of RpoB, which began to saturate at ∼50 µM RPP. Addition of rifampin blocked the labeling of RNAP by RPP, indicating that our probe is specific for the rifamycin-binding pocket (**Figure 4D**). If our hypothesis is correct and HelRSv can block/displace rifamycins, it should also affect labeling by RPP. We exposed RNAP to a gradient of HelRSv from 1:8 (HelRSv to RNAP) in the presence and absence of 1 mM ATP (**Figure 4E)**. At concentrations of HelRSv approaching a 1:1 molar ratio with RNAP, there is an ATP independent loss of RPP labeling; however, in the presence of ATP labeling is significantly lower at comparable HelR concentrations. These data demonstrate that HelRSv prevents rifamycin binding to RNAP, an activity stimulated by the presence of ATP.

**Figure 4.**
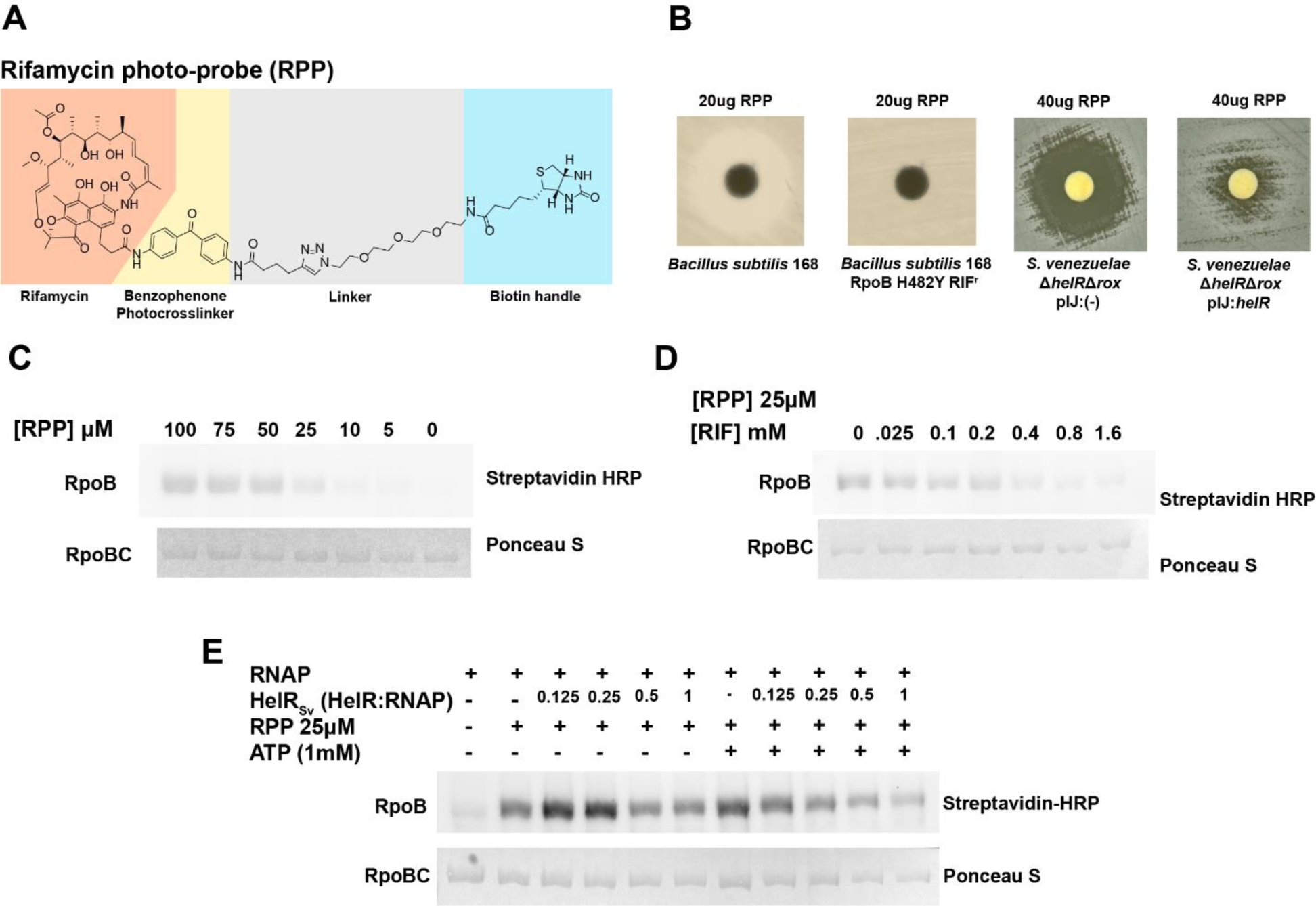
Design and activity of a rifamycin photoaffinity probe (RPP) demonstrate that HelRSv displaces RNAP-bound rifamycins. **A)** Structure of RPP with relevant elements highlighted. Rifamycin B scaffold (orange), Benzophenone (yellow), and Biotin (light blue) with the linker region in grey. **B)** Zones of growth inhibition resulting from RPP when spotted on a lawn of *B. subtilis* 168 wildtype and a rifampin resistant mutant (left) and *S. venezuelae ΔhelRΔrox* with empty vector or constitutive expression of *helR.* **C)** RPP labels RNAP *in-vitro*. RNAP was incubated with an increasing gradient of RPP, crosslinks were formed by exposure to 365nm light, after which proteins were analyzed by Western blot using Streptavidin-HRP for detection of covalently linked RPP followed by Ponceau S staining for total protein. **D)** Competition between Rifampin and RPP. Rifampin was added at various concentrations prior to addition of RPP (kept constant at 25µM). Samples were analyzed in the same manner as (A). **E)** HelRSv displaces RPP in an ATP-dependent manner. 25µM RPP was used to label RNAP which had been pre-incubated with HelRSv at various molar ratios, both in the presence and absence of 1 mM ATP.

### HelD-like proteins in microbial genomes

HelR belongs to a family of proteins not previously associated with antibiotic resistance. We surveyed the diversity of HelD-like proteins in bacteria and used the presence of a RAE to identify proteins likely involved in rifamycin resistance (HelRs). We ran BLASTp using HelRSv and HelDBs as queries and clustered the top 5000 hits at 99% identity, producing 4906 and 3026 representative sequences for HelR and HelD, respectively. As the non-redundant database includes sequences with varying levels of annotation, we used these sequences as a large BLASTp query set against consistently annotated genomes in the RefSeq database. Using the representative non- redundant HelD/HelR protein sequences to query RefSeq Actinobacteria and Firmicute genomes returned 15,136 putative HelD/HelR sequences. At a 50% identity threshold, the sequences can be assembled into 417 cluster families. We removed several spurious clusters of glycosyltransferases (see **Supplementary Table 2**, **Figure S5**) and then identified protein clusters with at least one member associated with a RAE. Only 29 of these cluster families, encompassing 3314 HelD-like sequences (21.9% of the total), contain at least one member associated with a RAE, and none of these originate from Firmicute genomes. 1774 protein sequences (out of 3314 or 53.5% of the sequences in the 29 RAE-associated clusters, 11.7% of the total putative HelD/HelR sequences) are associated with a RAE. 12 of these 29 clusters have only one member, and 21 clusters had <8 members, **Figure 5** shows a summary of the clusters with >8 members. We designated proteins as putative HelRs if they fell into a cluster with a high proportion of RAE association. The *S. venezuelae* and mycobacterial HelRs fall into the two largest clusters (235 and 74, respectively). Except for clusters 195 and 230, most proteins belonging to one of these clusters are associated with a RAE. This was notably lower in cluster 74, which encompasses the *Mycobacterial* sequences. We attempted to assemble all the HelD-like proteins into a comprehensive phylogeny to determine where the rifamycin resistance proteins fell on the tree, but these sequences proved too diverse. For instance, just the RAE-associated clusters 53, 68, 74, 124, 133, 195, 230, 235, and 266 produced an alignment with over 4000 sites (more than 4x the length of an individual HelD-like protein) and many gaps. Automated trimming reduces this alignment to 357 sites with visually poor alignment. We concluded that it was impossible to produce a reliable phylogeny using all these sequences; they are too diverse for the usual approaches. An alternative method to represent the relationships between these sequences is as a sequence similarity network (SSN)(Copp et al., 2018). We constructed an SSN from an all-against-all similarity search using an e-value cutoff of 1e^-10^. This network consists of 15137 nodes and 259,167 undirected edges after the removal of self-edges. We visualize this network in Cytoscape (Shannon et al., 2003) using the organic layout and include the cluster data from **Supplementary Table 2** (**Figure 5**). This network shows two distinct large clusters of HelRs (Clusters 235, 266, 68, 53) & (74, 230) and three small clusters. One consisting of all the members of 133, and two clusters made up of subsets of cluster 235 (**Supplementary figure 6**). We found no instances of Firmicute and Actinobacterial HelD-like proteins within a cluster. HelRs make up a distinct subset of all HelD-like enzymes, suggesting that most of these proteins are not involved in rifamycin resistance and may have analogous biological functions to *B. subtilis* HelD or perhaps novel ones.

**Figure 5.**
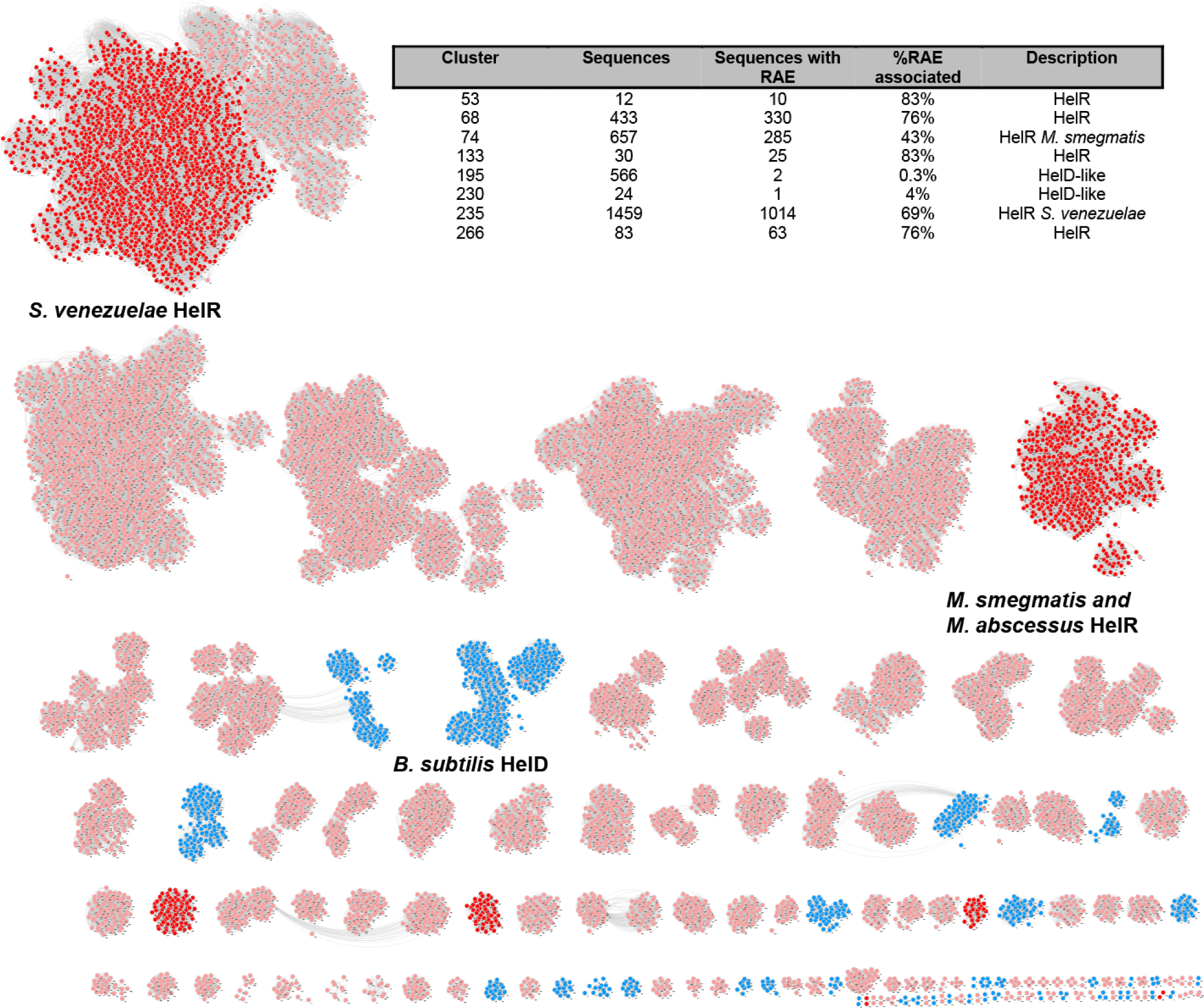
Sequence Similarity network of HelD-like proteins. A network of 15,137 HelD-like proteins from Refseq genomes of Actinobacteria and Firmicutes. Each protein is represented as a node, color coded by the phyla it comes from (light red for Actinobacteria, blue for Firmicutes) and whether or not it belongs to a protein cluster which is RAE associated (Red). Cluster 195 was deliberately not colored red because it contains so few RAE associated genes. HelRs (RAE associated HelD-like proteins) form two major clusters and 3 small ones, indicating most HelD-like proteins are not rifamycin resistance enzymes and likely have other functions.

**Figure 6.**
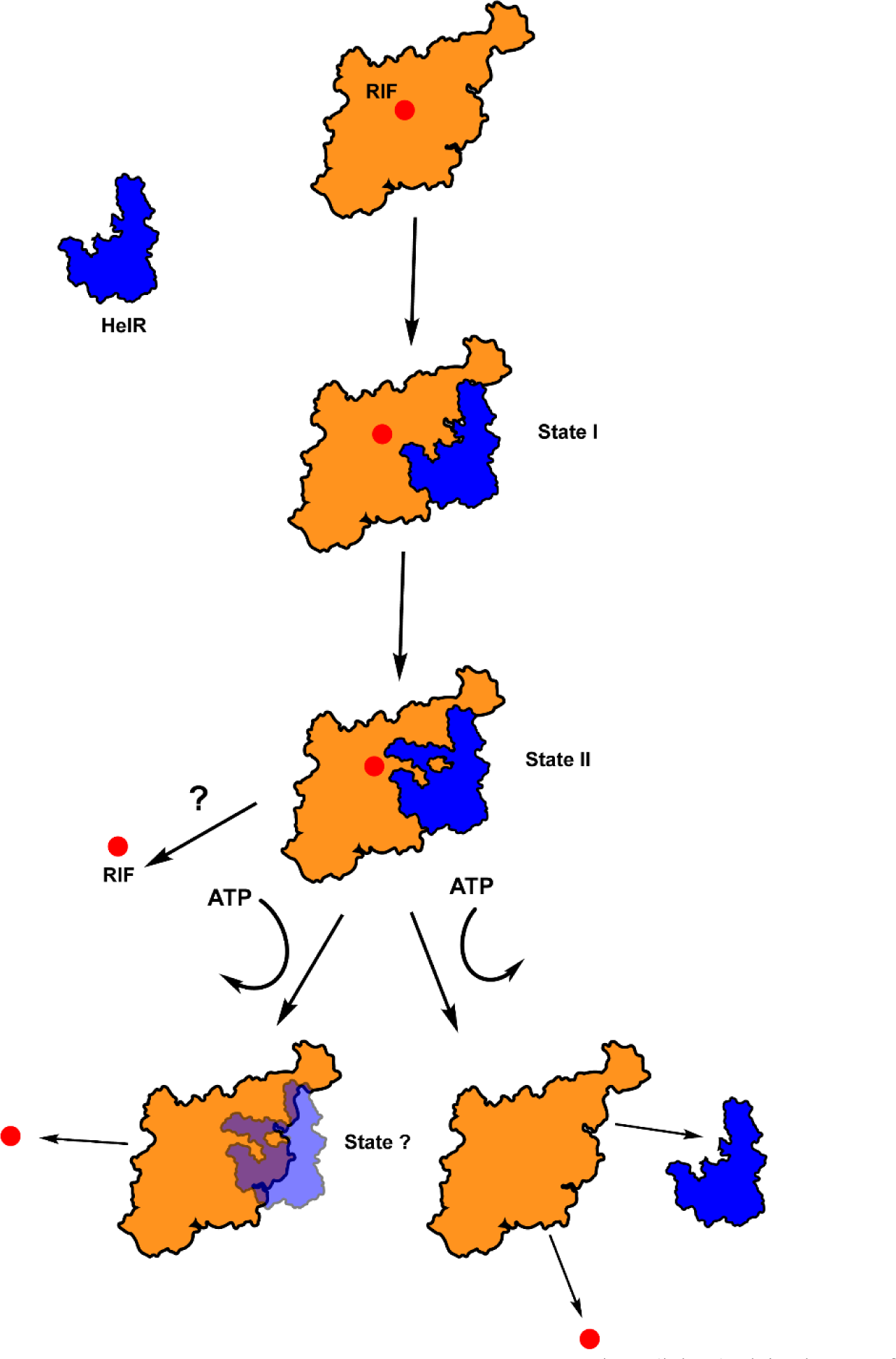
Model of HelR-mediated rifamycin resistance. HelR (blue) binds to free RNAP (orange) with rifamycin bound (RIF, Red circle). HelR isomerizes from State I to State II which may have a minor effect of rifamycin binding. We hypothesize that ATP hydrolysis occurs following the formation of a complex in State II and that the resulting conformational change leads to the active displacement of rifamycins. HelR may remain bound to RNAP following ATP hydrolysis or as in the case of *B. subtilis* HelD ATP hydrolysis may drive the dissociation of RNAP and HelR, this ambiguity is noted in the figure.

We also noted an abundance of HelD-like proteins in *Streptomyces* genomes. This is not solely due to the overrepresentation of *Streptomyces* genomes in Refseq and reflects numerous copies of HelD-like genes per genome. For instance, *S. venezuelae* ATCC 10712 encodes 5 HelD-like proteins (including HelRSv), whereas *M. smegmatis* and *M. abscessus* have only HelR and no other HelD-like proteins. HelR is also not strictly conserved among mycobacteria and is absent from medically important slow-growing species such as *M. tuberculosis*, M. *kansasii, M. ulcerans, M. leprae,* and *M. bovis*.

We were intrigued by cluster 195, which has 566 members, of which only two are associated with a RAE (**Figure 5**). *S. venezuelae* encodes a member of cluster 195, SVEN5092, which we identified in our RNAP preparations, evidence that this protein also interacts with RNAP. Unlike HelRSv, SVEN5092’s abundance was unchanged between rifamycin-induced and uninduced samples. We deleted this gene and were surprised to find that *S. venezuelae* Δ*5092* had become 4-fold *more* resistant to rifampin. When we performed the same deletion in a Δ*helR* background, resistance to rifampin was unchanged, suggesting that SVEN5092 may compete with HelRSv for RNAP binding. Nevertheless, SVEN5092 is an example of a HelD-like protein that does not confer rifamycin resistance. Curiously, even though it lacks a RAE, *helD* has been reported to be inducible by transcription inhibitors such as rifampin in *B. subtilis* (Hutter et al., 2004). This induction has been used as a reporter to identify the mechanism of action of novel antimicrobials (Mosaei et al., 2018). We confirmed that *B. subtilis* Δ*helD* is no more susceptible to rifampin than the parent strain. HelRs are a minority of HelD-like enzymes and are unique in their ability to confer rifamycin resistance.

## Discussion

The environmental resistome is the source of many of the antibiotic resistance elements that emerge in pathogenic bacteria (Surette and Wright, 2017). A complete understanding of the mechanistic diversity of antibiotic resistance is needed to preserve antibiotics in the resistance era (Brown and Wright, 2016). Characterizing these mechanisms can help us anticipate resistance before it emerges in the clinic, identify genes responsible for intrinsic resistance in pathogens, and guide the synthesis of new drugs. The capacity of bacteria to engage in horizontal gene transfer means that resistance in the environment remains an existential threat to the continued clinical efficacy of antibiotics over the long term.

In this work, we use the presence of a *cis-*regulatory genetic element that specifically induces genes in response to rifamycin antibiotics to guide the discovery of a novel resistance gene, *helR.* Using *S. venezuelae* as a model system, we confirm that the expression of *helR* is rifamycin inducible and that *helR* confers resistance to a variety of rifamycins, both natural and semisynthetic drugs in current use.

Based on a shared domain architecture with the known RNAP interacting protein HelD, we hypothesized and verified that HelRSv directly interacts with RNAP. HelD-like enzymes, HelR included, share the core ATPase machinery of Superfamily 1 helicases. These proteins use the energy released from ATP hydrolysis to generate mechanical force to promote translocation along ssDNA and strand displacement. In contrast to true helicases, HelD-like enzymes lack DNA binding domains and instead use this mechanical force to remodel themselves and/or their interaction partner RNAP (Kouba et al., 2020; Newing et al., 2020; Pei et al., 2020). We demonstrated that ATP hydrolysis is essential for the resistance activity of HelRSv but that HelRSv does not inactivate the antibiotic. This observation is analogous to another group of antibiotic resistance enzymes, the ribosomal protection proteins such as TetO/TetM. These proteins are GTPases that bind to the ribosome and directly displace target-bound tetracyclines, allowing for resumption of translation (Dönhöfer et al., 2012; Li et al., 2013). HelRSv is an ATPase which binds to RNAP, so we hypothesized that it directly displaces rifamycins, functioning as an RNAP protection protein. We directly tested this hypothesis *in vitro* and found that HelRSv does rescue transcription in inhibitory concentrations of rifamycins but by less than one order of magnitude.

This comes close to but does not fully recapitulate the fold change in MIC this gene confers (8-16 fold). We note that experiments with a HelR and RNAP from *Mycobacterium abscessus* show a comparable result in the same assay (P. Ghosh, personal communication). This could be a function of our *in vitro* system, which could be missing important factors. HelD, for instance, acts synergistically with the Firmicute specific δ subunit of RNAP (Pei et al., 2020; Wiedermannovà et al., 2014), perhaps there are other RNAP interaction partners required to capture the full extent of resistance *in vitro*. Data from HelD suggests that ATP hydrolysis favors dissociation from RNAP; if this is the case for HelR, it could give rifamycins a chance to re-bind RNAP before transcription can proceed to elongation masking the effect of HelR on free-RNAP *in vitro*. Rifamycin-bound RNAP also engages in abortive transcription, during which it occupies the template DNA’s promoter. These complexes would likely prevent free-RNAP from binding, also masking the effects of HelR on free-RNAP *in vitro*. Nevertheless, both the in cell and *in vitro* experiments are fully consistent with HelR’s importance in rifamycin resistance. It is also relevant to note that we do not observe any stimulation of transcription by HelRSv, offering further evidence that it is not functionally equivalent to HelD and instead has a dedicated function in antibiotic resistance.

To explore the displacement of rifamycins from the active site, we synthesized a photo-crosslinking rifamycin (RPP). Experiments with this probe revealed that HelRSv could indeed displace rifamycins from RNAP, confirming our hypothesis that HelRSv functions as a protection protein. Structural data from HelRMs implies that HelR and DNA cannot co-exist within RNAP, suggesting that HelR can only remove rifamycins from free-RNAP (Kouba et al., 2020; Newing et al., 2020; Pei et al., 2020). Incubation of HelRSv with 1mM ATP significantly reduced labeling by RPP compared to an incubation lacking ATP, suggesting that ATP hydrolysis may drive the displacement of rifamycins. Ribosomal protection proteins such as TetO and TetM are GTPases that use GTP hydrolysis to dissociate from the ribosome, not to directly displace tetracyclines. Their ability to displace tetracyclines is due to conformational changes they induce in the ribosome (Dönhöfer et al., 2012; Li et al., 2013). Dissociation of HelD from RNAP following addition of ATP has been reported by Pei *et al*. however if this were strictly required for dissociation, one would expect ATPase null HelD to be a potent inhibitor of transcription *in vitro,* but this is not the case (Newing et al., 2020). This fact, taken alongside our data that show that a combination of HelRSv and ATP more efficiently blocks binding of RPP to RNAP, suggests that conformational changes as a result of ATPase activity play a more critical role in HelR than it does for ribosomal protection proteins.

These data support a molecular model of HelR-mediated resistance (**Figure 6**). Following exposure to rifamycin antibiotics, HelR is produced and associates with RNAP. The structures of HelRMs suggest that this begins with HelR in state I, with a disordered PCh loop, which subsequently folds into the primary channel generating state II. We hypothesize that this state directly precedes ATP binding and hydrolysis. Conformational changes driven by ATP hydrolysis are then responsible for displacing rifamycins from their binding pocket. The exact nature of these conformational changes is currently unknown but likely also results in the dissociation of HelR from RNAP. Rifamycins cannot inhibit RNAP once transcription has begun; by constantly removing these antibiotics from free-RNAP, HelR maintains a pool of transcriptionally-competent RNAP, thereby conferring resistance. In support of this mechanism, Kouba *et al*. show that HelRMs can form a complex with RNAP with both σ^A^ and RpbA bound, facilitating transcription initiation following removal of rifamycins (Kouba et al., 2020).

Microbes exhibit tolerance to antibiotics when they survive a transient exposure to inhibitory levels of a bactericidal antibiotic (Brauner et al., 2016). Rifamycins show bactericidal activity against Gram-positive organisms such as the mycobacteria (Floss and Yu, 2005), making this drug class highly important for treating tuberculosis. Despite this, the mechanism by which rifamycins kill bacteria is not well understood. It has been tied to aberrant central metabolism and reactive oxygen species generation in multiple studies (Lobritz et al., 2015; Piccaro et al., 2014). Several known mechanisms of tolerance have been characterized, including rifampin efflux pumps, overproduction of RNA polymerase, and even mistranslation of RpoB, leading to the generation of a subpopulation of rifamycin-resistant RNAP (Adams et al., 2011; Javid et al., 2014; Zhu et al., 2018). In this study, we demonstrate that deletion of *helR* decreases tolerance of *S. venezuelae* to rifampin. An effect that is not observed for the rifamycin inactivating enzyme Rox. HelR’s ability to displace rifamycins from RNAP likely sustains some level of transcription in the presence of high concentrations of rifamycins. We note that rifamycin-mediated cell death is rapid and that many bacterial strains with a RAE-associated *helR* also have RAE-associated genes encoding rifamycin inactivating enzymes. We speculate that HelR’s principal role may be to provide rifamycin tolerance following drug exposure to enable sufficient destruction of the antibiotic in the local environment by RAE-associated inactivation enzymes. This hypothesis would explain why many bacterial strains have both HelR and rifamycin inactivating enzymes under control of RAEs – HelR’s provide tolerance in the presence of antibiotic, offering sufficient time for inactivating enzymes to decrease the concentration of drugs below the MIC.

HelD-like enzymes are numerous in the Firmicute and Actinobacteria phyla. We show that a small fraction of these are associated with RAEs and, therefore, likely to contribute to rifamycin resistance. Much of the diversity within this large family of proteins has yet to be interrogated. Some may be involved in resistance to other RNAP inhibitors, or they may function to aid in transcriptional cycling like HelD. Of the fraction that are involved in resistance, many copies are found in pathogenic bacteria. One particularly relevant example is *M. absc*essus (P. Ghosh, personal communication), but HelRs are present in many fast-growing mycobacteria. These genes are also found in pathogenic *Nocardia* species such as *N. farcinia* and the foal pathogen *Rhodococcus equi*. The identification and understanding of HelR can now guide the synthesis of rifamycin analogs that are not substrates for HelR, leading to more effective rifamycin chemotherapy for various infections in the resistance era.

## Acknowledgments

We thank P. Ghosh for sharing preliminary data on HelR from *M. abscessus*. This work was funded by the Canadian Institutes of Health Research (FRN-148463) and by a Canada Research Chair to G.D.W. M.D.S. is supported by a Graduate Scholarship from the Natural Sciences and Engineering Council of Canada.

## Author contributions

M.D.S Conceived and performed experiments, wrote the manuscript. K.K and N.W. Performed experiments and provided expertise and feedback. G.D.W conceived of experiments, wrote and edited the manuscript.

## Methods

### Bacterial Strains and Conditions

All bacterial species and strains used in this study are listed below *S. venezuelae* ATCC 10712 was grown using Bennett’s media (10 g potato starch, 2 g casamino acids, 1.8 g yeast extract, and 2 mL of Czapek’s mineral mix [10 g KCl, 10 g MgSO4⋅7H2O, 12 g NaNO3, 0.2 g FeSO4, 200 μL concentrated HCl in 100 mL H2O] per 1 L H2O) or Tryptic Soy Broth (BD Bacto^TM^) with and without 1.5% agar depending on the application. Biparental matings took place on Soy Flour Mannitol (20 g soy flour, 20 g mannitol and 20 g agar) plates supplemented with 15 mM MgCl2. *Escherichia coli* was grown using L.B. (Lysogeny broth) except where specified. *B. subtilis* was grown using Tryptic Soy Broth (BD Bacto^TM^). Antibiotics for selection were used at the following concentrations 7.5 µg/mL kanamycin, 100 µg/mL ampicillin, 34 µg/mL chloramphenicol, 50 µg/mL apramycin, 100 µg/mL and 50 µg/mL hygromycin B for *E. coli* and *S. venezuelae,* respectively.

### Strains and plasmids used in this study

| Strain or Plasmid | Genotype/Use | Source or Reference |
| --- | --- | --- |
| *Streptomyces venezuelae* |  |  |
| ATCC 10712 |  | ATCC |
| $\Delta helR$ | $\Delta SVEN\_6029(helR)::[aac(3)IV-oriT]$ | This study |
| $\Delta rox$ | $\Delta SVEN\_0481(rox)::[aac(3)IV-oriT]$ | This study |
| $\Delta helR-scar$ | $\Delta SVEN\_6029(helR)::[FLP scar]$ | This study |
| $\Delta helR$ pIJ10257 | $\Delta SVEN\_6029(helR)::[aac(3)IV-oriT]$<br>φBT1::pIJ10257 (empty) | This study |
| $\Delta helR$ pIJ10257:*helR* | $\Delta SVEN\_6029(helR)::[aac(3)IV-oriT]$ φBT1::pIJ10257: *helR* | This study |
| $\Delta helR\Delta rox$ | $\Delta SVEN\_6029(helR)::[scar]$<br>+ (*rox*)::[*aac(3)IV-oriT*] | This study |
| $\Delta helR\Delta rox$<br>pIJ10257 | $\Delta SVEN\_6029(helR)::[scar]$<br>+ (*rox*)::[*aac(3)IV-oriT*]<br>φBT1::pIJ10257 (empty) | This study |
| $\Delta helR\Delta rox$<br>pIJ10257:*helR* | $\Delta SVEN\_6029(helR)::[scar]$<br>+ (*rox*)::[*aac(3)IV-oriT*]<br>φBT1::pIJ10257: *helR* | This study |
| $\Delta helR \Delta rox$<br>pIJ10257:*helR*<sub>D533A</sub> | $\Delta SVEN\_6029(helR)::[scar]$<br>+ (*rox*)::[*aac(3)IV-oriT*]<br>$\phi BT1::pIJ10257: helR_{D533A}$ | This study |
| pGUS:*P<sub>rox</sub>* | $\phi C31::pGUS:P_{rox}$ | This study |
| $\Delta helR$ pGUS:*P<sub>rox</sub>* | $\Delta SVEN\_6029(helR)::[aac(3)IV-oriT]$ $\phi C31::pGUS:P_{rox}$ | This study |
| $\Delta rox$ pGUS:*P<sub>rox</sub>* | $\Delta SVEN\_0481(rox)::[aac(3)IV-oriT]$ $\phi C31::pGUS:P_{rox}$ | This study |
| $\Delta helR \Delta rox$ pGUS:*P<sub>rox</sub>* | $\Delta SVEN\_6029(helR)::[scar]$<br>+ (*rox*)::[*aac(3)IV-oriT*]<br>$\phi C31::pGUS:P_{rox}$ | This study |
| $\Delta 5092$ | $\Delta SVEN\_5092::[aac(3)IV-oriT]$ | This study |
| $\Delta helR \Delta 5092$ | $\Delta SVEN\_6029(helR)::[scar]$<br>+ $\Delta SVEN\_5092::[aac(3)IV-oriT]$ | This study |
| **Bacillus subtilis** |  |  |
| Strain 168 |  | ATCC 23857 |
| Rif resistant |  | (Spanogiannopoulos, 2014) |
| $\Delta helD$ | *helD(yvgS)::[kan]* | Gift from Eric Brown. Construct is from the *B. subtilis* strain collection (Koo et al., 2017) |
| **Amycolatopsis mediterranei** |  |  |
| Strain S699 |  | Gift from Tin-Wein Yu |
| **Escherichia coli** |  |  |
| DH5α | General cloning | Invitrogen |
| ET12567/pUZ8002::*bla* | Chlor<sup>r</sup>; Amp<sup>r</sup>; Methylation deficient strain used for conjugation into *Streptomyces* | (MacNeil et al., 1992) |
| DH5α pCP20 | Amp<sup>r</sup>; FLP mediated recombination | (Cherepanov and Wackernagel, 1995) |
| BW25113 pKD46 | Amp<sup>r</sup>; $\lambda$ red recombineering | (Datsenko and Wanner, 2000) |
| **Plasmids or Cosmids** |  |  |
| pIJ10257 | Hyg<sup>r</sup>; $\phi BT1$ integrative vector for protein expression from the strong constitutive *ermE* promoter. | (Hong et al., 2005) |
| pIJ:*helR<sub>sv</sub>* | *SVEN_6029(helR)* cloned into pIJ10257 using NdeI/HindIII sites | This study |
| pIJ:*helR<sub>sv</sub>* D538A | *helR* with a D538A substitution cloned into pIJ10257 using NdeI/HindIII sites | This study |
| pGUS | Apra<sup>r</sup> Spec<sup>r</sup>; $\phi C31$ integrative vector containing promoterless *gusA* | (Myronovskyi et al., 2011) |
| pGUS:*P<sub>rox</sub>* | *rox* promoter cloned into KpnI/XbaI sites on pGUS | This study |
| SV-2 G10 | Supercos-1 cosmid containing *SVEN_0481 (rox)* | Gift from MJ Buttner |
| SV-2 C08 | Supercos-1 cosmid containing *SVEN_6029 (helR)* | Gift from MJ Buttner |
| SV-5 E02 | Supercos-1 cosmid containing *SVEN_5092* | Gift from MJ Buttner |

### RNA isolation

Approximately 10^9^ spores of *Streptomyces venezuelae* ATCC 10712 were used to inoculate a 50 mL starter culture in Bennett’s media. Following incubation for 16h at 30°C with shaking (250 rpm), this culture was used to seed 6 x 200 mL subcultures (with a 1:100 dilution of starter). These subcultures represent two sets of biological triplicates, one receiving rifamycin SV treatment (0.5 µg/mL) and vehicle control (DMSO). After 8 hours of growth the cultures were treated, and returned to the incubator for another two hours of growth after which 10 mL of culture were collected in a 15 mL Falcon tube and centrifuged for 10 minutes at 8 000 x g at 4°C. The supernatant was discarded, and the pellets were flash frozen in liquid nitrogen and stored at -80°C until RNA isolation.

Prior to RNA isolation, cell pellets were incubated with 300 µL of 10 mg/mL lysozyme (prepared fresh in sterile H2O) and incubated for 7 minutes at 30°C/250 rpm to allow for efficient lysis by TRIzol reagent (Ambion). 10-15 glass beads (4mm diameter) and 4 mL of TRIzol reagent were added to each cell pellet. Samples were then vortexed for 5 minutes and 0.8 mL of chloroform was added followed by 4 cycles of 30 seconds vortexing followed by 30 seconds on ice. The glass beads were removed, and the phenol/chloroform mixture was separated using centrifugation (10 minutes at 8 000 x g and 4°C). The aqueous phase was removed and mixed with an equal volume of phenol-chloroform (pH 4, Ambion). This solution was vortexed for 30s and centrifuged once more. The aqueous phase was mixed with an equal volume of 75% ethanol and applied directly to a column from the Purelink^TM^ RNA Mini Kit (Invitrogen) and the rest of the purification is performed according to the manufacturer’s instructions. RNA integrity was qualitatively examined by gel electrophoresis. cDNA was synthesized from 1 µg of RNA using the Maxima First Strand cDNA synthesis kit (Thermo Scientific) according to the manufacturer’s recommendations for high GC content templates.

### RT-qPCR

Quantitative PCR was performed using SYBR Select Master Mix CFX (Applied Sciences), all primer concentrations were 200 nM, cDNA was diluted 1:1 with nuclease free water (Ambion) and 2 µL of diluted cDNA was used as template. Thermocycling conditions were as follows, 2 minutes at 50 °C, 2 minutes at 95 ° C, then 40 cycles of (95 °C for 15s, 57 °C for 15s, and 72 °C for 30s). Quantitative PCR was performed on a Bio-Rad C1000 Thermocycler. Gene specific primers are as follows *hrdB*-q F.P. and R.P., *rox*-q F.P. and R.P., *helR-q F.P.* and R.P. Expression is reported as a ratio of the housekeeping gene *hrdB.* P-values were determined using an unpaired students t-test.

### Mutant construction

Deletions of *rox* (*SVEN_0481*), *helR*(*SVEN_6029*) and *SVEN5092* were made using the REDIRECT method^19^. PCR targeting of cosmids SV-2 G10 (containing *rox)*, SV-2 C08 (containing *helR*), and SV-5 E02 (containing SVEN5092) using primer pairs *rox* K.O. FP/RP, *helR* K.O. FP/RP, and *SVEN5092 K.O. FP/RP* generated in frame deletions of all genes by replacing them with an *oriT-*apramycin resistance cassette. Cosmids were introduced to *S. venezuelae* by biparental mating. Successful recombination was identified by isolates with Apra^r^/Kan^s^ sensitivity profile and confirmed using amplification primers flanking the deletions (*rox* cPCR FP/RP, *helR* cPCR FP/RP, *SVEN5092 cPCR FP/RP*). *S. venezuelae* Δ*helR*Δ*rox* was constructed by removing the *oriT* apramycin cassette from mutagenized SV-2 C08 using FLP recombinase. An *oriT* hygromycin B resistance cassette was then inserted into the cosmid backbone using PCR targeting. This cosmid was introduced into *S. venezuelae* Δ*helR* by conjugation and selecting for single recombination (Hyg^r^). Individual colonies were screened until a double recombinant could be found (Apra^s^/Hyg^s^), to create *S. venezuelae* Δ*helR-scar.* The removal of the *oriT* apramycin cassette was confirmed by PCR, *rox* was subsequently knocked out in *S. venezuelae* Δ*helR-scar* using mutagenized SV-2 G10 as described above, yielding *S. venezuelae* Δ*helR*Δ*rox. S. venezuelae* Δ*helR*-scar was also used to construct *S. venezuelae* Δ*helR*Δ*5092*.

### Antibiotic susceptibility testing

Antimicrobial susceptibility testing was performed according to general broth microdilution CLSI protocols^53^. *S. venezuelae* inoculum for MICs was grown in 3mL TSB with 2-3 4mm glass beads for 48 hours at 30°C with shaking 250rpm. Under these conditions *S. venezuelae* does not form large clumps/aggregates and cultures can be reliably standardized by OD600nm. All *S. venezuelae* MICs were determined in TSB following incubation at 30°C for three days with shaking (250rpm).

### Antibiotic inactivation assay

*S. venezuelae* spores were used to inoculate 6 mL of Tryptic soy broth (TSB) which was incubated at 30 °C for 24 hours with shaking (250rpm). The cells were then collected by centrifugation (5 000 x g for 10 minutes) and then resuspended in 3mL of TSB + 5 µg/mL rifamycin SV and incubated for another 24 hours. These cultures were centrifuged once more and 20µL of supernatant was added to cellulose discs and placed on top of a lawn of rifamycin susceptible *B. subtilis* 168 grown on TSA. *B. subtilis* was incubated at 30 °C for 16 hours and then imaged.

### Antibiotic time kill assay

Spores of *S. venezuelae* were cultured overnight in TSB, then diluted 1:100 and allowed to grow for 6 hours to reach exponential phase. These cultures were standardized to an OD600nm of 0.1 and rifampin (10X MIC of each individual strain) or DMSO was added. At each of the specified timepoints (0h, 8h and 24h) 1mL of culture was harvested by centrifugation (10 000 x g 3 minutes) and resuspended in 1mL of fresh TSB. Centrifugation and resuspension were repeated two additional times to remove all residual rifampin. Serial 10-fold dilutions of culture were then spotted onto Bennett’s agar and incubated for 24 hours before imaging. Bennett’s was chosen over TSB for this application because *S. venezuelae* forms large amorphous colonies on TSA making these spot dilutions difficult to interpret.

### *S. venezuelae* RNA polymerase purification

Native RNA polymerase was purified from *S. venezuelae* using the procedure outlined in(Kieser et al., 2000), based on the protocol from Burgess and Jendrisak (Burgess and Jendrisak, 1975). With the following modifications: HEPES was substituted for Tris in all buffers and a HiPrep Heparin FF 16/10 column (Cytiva) was used in place of a DNA-cellulose column. A Superdex 200 10/300 GL column (G.E. Life Sciences) was used for gel filtration. For *in-vitro* transcription and labelling experiments RNAP was further purified using Capto HiRes Q (Cytiva). RNAP containing fractions were pooled and concentrated using Amicon Ultracentrifugal filters (EMD Millipore). Concentrated protein was diluted 1:1 with glycerol and stored at -20 until use. The same growth conditions established for RNA isolation were used to grow log phase *S. venezuelae* for proteomics experiments. For bulk isolation of RNA polymerase (for *in-vitro* transcription and RPP labelling) *S. venezuelae* Δ*helR* was grown for 24 hours in TSB as this yielded more biomass than Bennett’s media.

### Co-Immunoprecipitation assays

*S. venezuelae* Δ*helR*Δ*rox* pIJ:*helR* and pIJ:*helR*-FLAG spores were used to inoculate 3mL of TSB cultures. The next day these cultures were used to inoculate 50 mL of fresh TSB media which was incubated for 16h before the cells were harvested by centrifugation at 10 000 x g for 10 minutes. Cells were resuspended in Tris-buffered saline (TBS) with lysozyme (Bioshop), DNase (Bovine Pancreas, Sigma Aldrich), and 1 Peirce Protease Inhibitor tablet (Thermo Fisher) and subsequently lysed by two passages through a cell disruptor (Constant Systems Limited, Daventry U.K.) at 30k PSI. Insoluble protein was removed by centrifugation at 30 000 x g for 30 minutes at 4 °C. Soluble protein was quantified using a Pierce^TM^ BCA Protein Assay Kit (ThermoFisher) and standardized to 2 mg/mL. 1mL (2 mg) of protein was added to 20 µL of ANTI-FLAG^®^ M1 Agarose affinity Gel (Sigma Aldrich) pre-equilibrated in TBS and incubated on a nutator for 2 hours at room temperature. Beads were collected by centrifugation (6 000 x g for 30 seconds), the supernatant was discarded, and the beads were washed with 1mL of fresh TBS. 4 additional washes of the beads were carried out as described. 60 µL of 300 µg/mL FLAG peptide (GenScript) was used to elute protein from the beads.

Eluted protein was separated on an 8% polyacrylamide gel and transferred onto PolyScreen PVDF transfer membrane (Perkin Elmer). Membranes were blocked overnight in TBS containing 5% Bovine Serum Albumin (Sigma Aldrich). After three 10-minute washes with TBST (0.1% tween 20) membranes were probed with a 1:5000 dilution of an α-FLAG HRP conjugate for detection of HelRSv-FLAG, or with a 1:5000 dilution of mouse α-RpoB (8RB13, Fisher Scientific) followed by a 1:20000 rabbit α-mouse HRP conjugate (Ab97046, Abcam) for detection of RNA polymerase. Western blots were imaged using SuperSignal^TM^ West Pico PLUS chemiluminescence detection reagents as per manufacturer’s instructions (Fisher Scientific).

### HrdB purification

*E. coli* BL21 (DE3) plysS pET28a:*hrdB* was grown in 4L L.B. at 37 °C until an OD600nm 0.6, at which point it was cooled to 16 °C and induced with 0.1mM IPTG. Following a 10 hour induction cells were collected by centrifugation (6 000 x g for 15 minutes at 4 °C). Cells were resuspended into lysis buffer (25 mM HEPES pH 7.4, 400 mM NaCl, 15 mM imidazole, 1 mM DTT) with a protease inhibitor tablet, lysozyme, and DNase. Cells were lysed on a cell disruptor (Constant Systems Limited, Daventry U.K.) at 20k PSI and soluble protein was isolated by centrifugation at 30 000 x g for 30 minutes at 4 °C. Soluble protein was mixed with Ni-NTA resin (Qiagen) for 1h at 4 degrees on a nutator. Resin was collected by packing into a column and extensively washed with lysis buffer until protein could not be detected in the flow through. HrdB was eluted from the column by successive washes with lysis buffer containing increasing concentrations of imidazole (50 mM, 100 mM, 200 mM, 300 mM). Fractions containing HrdB were pooled and dialyzed against lysis buffer with no imidazole. The His-tag was cleaved by overnight incubation with thrombin (from bovine plasma, Sigma Aldrich) at 4 °C during dialysis. Tag free HrdB was recovered by passage through a Ni-NTA column, concentrated, and purified further by gel filtration (Superdex 200 Increase 30/100 GL). Pure fractions were pooled, concentrated, snap frozen in liquid nitrogen and stored at -80 degrees C until use.

### HelRSv purification

4L of TSB were inoculated with 1:500 dilution of *S. venezuelae* Δ*helR* pIJ:*helR*-*his6* overnight culture and incubated at 30 °C for 16 hours with shaking (250 rpm). Cells were collected by centrifugation (8 000 x g for 15 minutes at 4 degrees). Lysis was carried out as described during Co-IP except cells were resuspended in Lysis buffer (25 mM HEPES pH 7.4, 400 mM NaCl, 15 mM imidazole, 1 mM DTT). Ni-NTA chromatography was performed in the same manner as for HrdB, fractions containing pure HelRSv were pooled and dialyzed against Lysis buffer lacking imidazole, then concentrated, snap frozen, and stored at -80 °C until use.

### Proteomics

Protein mass spectrometry and proteome abundance analysis were performed at the Proteomic Core Facility of the Institute for Research in Immunology and Cancer at the Université de Montréal.

### *In-vitro* transcription

Multiple round transcription assays were performed in transcription buffer (40 mM Tris pH 7.9, 10mM MgCl2, 1.5 mM DTT, 0.25 mg/mL BSA, 20% (v/v) glycerol). Reactions (15 µL volume) were performed by incubating *S. venezuelae* RNA polymerase (25 nM) with σ^HrdB^ (100 nM) for 5 minutes at 30 °C, followed by HelRSv and rifampin (in relevant reactions) with an additional 5- minute incubation. Template DNA (PermE*) was added to a final concentration of 10 nM and 2 µL of NTPs (0.4 mM ATP, GTP, UTP, and 0.05 mM CTP with 20 µCi[α-^32^P] CTP (800 Ci/mmol)) were added to initiate the reaction which was allowed to proceed for 10 minutes at 30 °C. Reactions were stopped by the addition of an equal volume of loading buffer (7 M urea, 0.01% bromophenol blue, 20% glycerol) and analyzed on a 6% polyacrylamide gel containing 7 M urea. Transcripts were visualized using autoradiography.

PermE* DNA was synthesized as a gBlock (**Supplementary Figure 2**) with flanking *Hind* III and *Bam* HI sites and cloned into pUC19. Sequence verified plasmid was used as template for PCR using PermE* F.P. and PermE* R.P. PCR reactions were analyzed for purity on an agarose gel and purified using GeneJET PCR purification kit (Thermo Fisher).

### Rifamycin B purification

*Amycolatopsis mediterranei* S699 was grown on solid Bennett’s media for 7 days at 30°C and 600 mL of YMG media (4 g glucose, 4 g yeast extract, 10 g malt extract in 1 L distilled water) was inoculated with ∼10 colonies worth of biomass. Following 7 days of growth (30 °C and 250 rpm), production of rifamycin B was confirmed by LC-ESI-MS. After verifying rifamycin B production, the culture was mixed with an equal volume of methanol and returned to the shaker for 5 minutes, cells and other particulate were removed by centrifugation (15 minutes, 6 000 x g) and the supernatant was decanted. Methanol was removed from the sample by rotary evaporation and the remaining solution was lyophilized. Dried extract was resuspended in a 1:1 acetonitrile: H2O mixture, centrifuged to remove insoluble material (8000 x g for 5 minutes) and loaded onto a CombiFlash ISCO (RediSep^®^RF C18 26g (Teledyne)) and purified by reverse phase flash chromatography using a water-acetonitrile linear gradient to elute rifamycins. Fractions containing high absorbance at 306 nm were checked for purity by LC-ESI-MS. Fractions containing pure rifamycin B were pooled and lyophilized.

### Synthesis of RPP

#### a) Synthesis of N-[4(4-aminobenzoyl)phenyl]hex-5-ynamide (BPh-alkyne)

This compound was synthesized according to previously published methods (Salisbury and Cravatt, 2008). 1 mL (8.9 mmol) of 5-hexynoic acid was added to a 50 mL round bottom flask followed by 15 mL DMF (anhydrous, Sigma Aldrich), 1.7 g (8.9 mmol) 1-ethyl-3-(3- dimethylaminopropyl)carbodiimide (EDAC, Sigma Aldrich), and 1.2 g (8.9 mmol) hydroxybenzotriazole (HOBt, Sigma-Aldrich). This solution was incubated for 5 minutes at room temperature with stirring. After the solution clarified, 1.9 g (8.9 mmol) 4,4’-diaminobenzophenone (Sigma-Aldrich) was added and the reaction (now a brown color) was incubated at room temperature for 24 hours with stirring. Reaction was applied to a reverse phase CombiFlash ISCO (RediSep®RF C18 26 g (Teledyne)) and purified using a water-acetonitrile linear gradient. Fractions were monitored by LC-ESI-MS and pure fractions were pooled and lyophilized.

#### **b)** Coupling of BPh-alkyne to rifamycin B to form RP-Alkyne

85 mg (0.1125 mmol) of rifamycin B was dissolved in 1 mL of DMF in a 50 mL Falcon tube. Once rifamycin B had dissolved, 21.6 mg (0.1125 mmol) EDAC and 15.2 mg (0.1125 mmol) HOBt were added. This solution was incubated at room temperature with stirring for 20 minutes and then 34.6 mg (0.1125 mmol) BPh-alkyne were added and the reaction was allowed to procced for 16 hours at room temperature. Monitoring of the reaction by LC-MS showed considerable starting product remaining after 16 hours so 64.8 mg (0.3375 mmol) EDAC were added and the reaction was incubated for another 32 hours (48 hours total) after which no starting material could be observed by LC-MS. Reaction contents were applied directly to a LH-20 size exclusion column, compounds were eluted with methanol and collected using a BioRad model 2110 fraction collector and analyzed by LC-ESI-MS. Fractions containing Rifamycin B-N-[4(4- aminobenzoyl)phenyl]hex-5-ynamide (RP-Alkyne) were further purified by reverse phase flash chromatography using methodology previously described for rifamycin B (RediSep®RF C18 26 g (Teledyne)).

#### **c)** Huisgen cycloaddition to produce RPP

4.5 mg (0.004 mmol) of RP-Alkyne was dissolved in 50 µL of t-butanol then added to 30 µL H2O and vortexed briefly. 2.2mg (0.005 mmol) of Azide-PEG3-biotin (Sigma-Aldrich) was suspended in 10 µL of DMSO and added to the t-butanol:H2O mixture, followed by 3 µL sodium ascorbate (250 mg/mL in H2O) and 3 µL of CuSO4 (50 mg/mL in H2O), vortexing after each addition. Huisgen cycloaddition reaction was allowed to proceed for 2 hours after which RPP was purified by reverse phase flash chromatography as described previously using a RediSep®RF C18 4.3g column (Teledyne). Full compound analysis by NMR and mass spectrometry is provided in Supplementary Figures 7-14, and Table S3.

### RPP Photolabeling

Photoabeling reactions were carried out at 15µL scale in TBS. *S. venezuelae* RNA polymerase, HelRSv, ATP were prepared as 15X stocks in TBS, and Rif and RPP were dissolved in DMSO as 15X stocks. 130 nM of RNA polymerase was incubated with HelRSv, ATP, and Rif where indicated for 5 minutes, after which RPP was added and allowed to incubate for another 5 minutes in total darkness. Parafilm was spread over a 96 well plate and a gloved hand was used to create indentations in each well. Labelling reactions were aliquoted into these indentations and the 96 well plate was placed under a UVP Blak-Ray B-100A lamp (365nm, Analytik Jena) approximately 20 cm from the bulb for 10 minutes. Reactions were mixed with 5uL of 4X Laemmli sample buffer (Bio-Rad), boiled for 5 minutes, separated on a 6% polyacrylamide gel, and transferred to PVDF membranes. These blots were blocked, washed, and imaged as described for α-FLAG and α-RpoB blots but probed using a 1:25000 dilution of Streptavidin-HRP conjugate (Sigma Aldrich).

### HelR/HelD network analysis

This pipeline is available at https://github.com/waglecn/helD_search and is written in Snakemake v6.0.5 (Köster and Rahmann, 2018) In order to ensure a minimum consistent annotation for targeted genomes, all RefSeq assemblies from organisms classified according to the NCBI taxonomy in the phyla Actinobacteria and Firmicutes were downloaded using the ncbi-genome-download v0.3.0 (https://github.com/kblin/ncbi-genome-download), totaling 7457 genomes as of Jan. 12^th^ 2021.Up to 500bp upstream of each coding sequence feature from each of the downloaded genomes was extracted into separate fasta files. The previously published set of RAE sequences was used to generate an alignment and nucleotide Hidden Markov Model (HMM) using nhmmer from the HMMER 3.1b software (Eddy, 2011; Spanogiannopoulos et al., 2014). This RAE HMM was used to search every upstream sequence of every CDS of every downloaded genome. The search was limited to the top strand only, and putative RAE sequences were identified as having a minimum length of 15 bp and a minimum score of 10.0 bits.

The HelR sequence from *M. smegmatis* (WP_003893549.1) was used to search the NCBI nr database using BLASTp with default parameters (Oyama et al., 2008). The top 5000 sequence hits (by E-value) were downloaded as a fasta file, sorted by length, and subjected a clustering step using usearch v11.0.667 using a threshold of 99% identity (Edgar, 2010). This resulted in a set of 4906 clusters. The centroid sequence from each cluster was used to query each of the downloaded genome assemblies using an E-value cutoff of 1e-255 using DIAMOND v 0.9.14.115 (Buchfink et al., 2014). This resulted in a set of 15,136 putative HelD sequences. These sequences were annotated with the RAE sequence, if identified, in their upstream sequences.

The RAE annotated HelD sequences from RefSeq Actinobacteria and Firmicutes were then sorted by length and subjected to a cluster analysis using usearch at a threshold of 50% identity, resulting in 161 clusters. 15 clusters were identified having at least one member sequence possessing an upstream RAE (Edgar, 2010).

The 3988 RAE-associated HelD (putative HelR) sequences were aligned using mafft v7.310 (using the “—auto” parameter) resulting in a highly gapped alignment of 3874 columns(Katoh and Standley, 2013). This alignment was trimmed using TrimAl v1.4.rev22 (using the “--automated1” parameter) to an alignment of 357 columns (Capella-Gutiérrez et al., 2009).

To generate the SSN, the combined set of sequences identified in the RefSeq genomes using the HelD search set were subjected to an all-vs-all search using DIAMOND (Buchfink et al., 2014) with an E-value cutoff of 1x10^-10^. Self-hits were filtered out of these results. The network was further refined by using a minimum edge score cutoff of 700 bits for each similarity result to produce the final network.

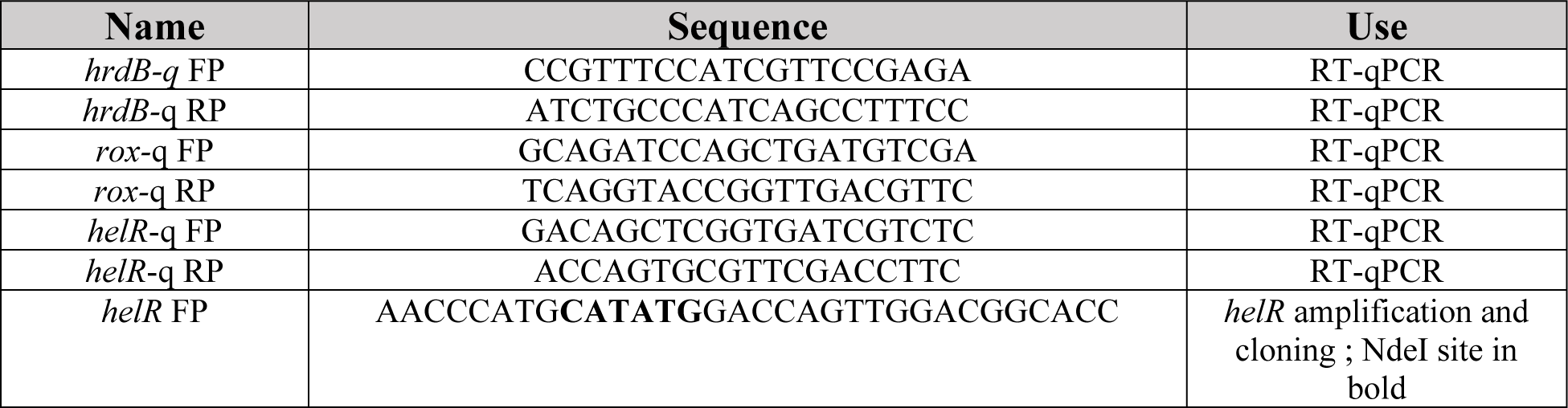
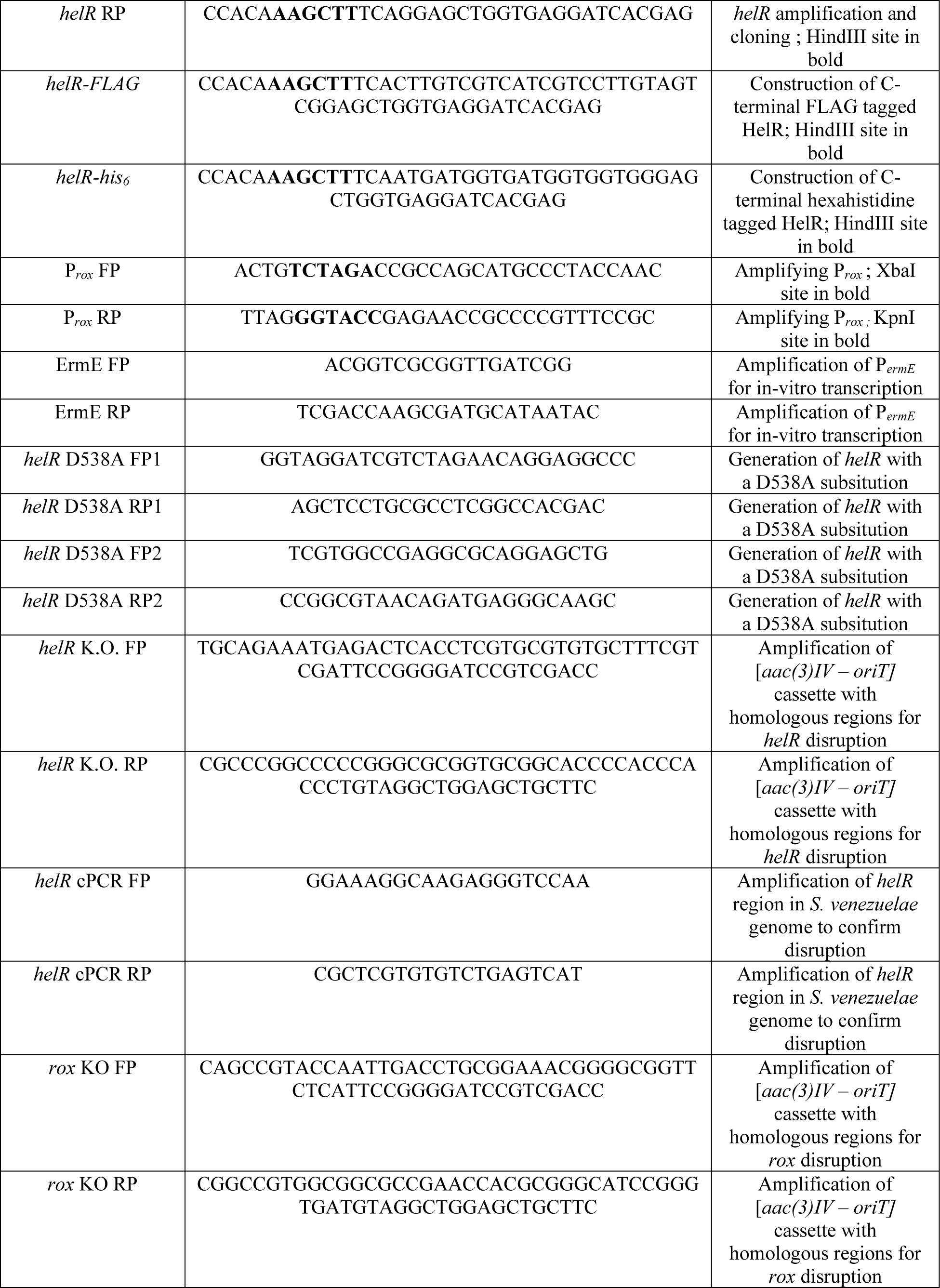
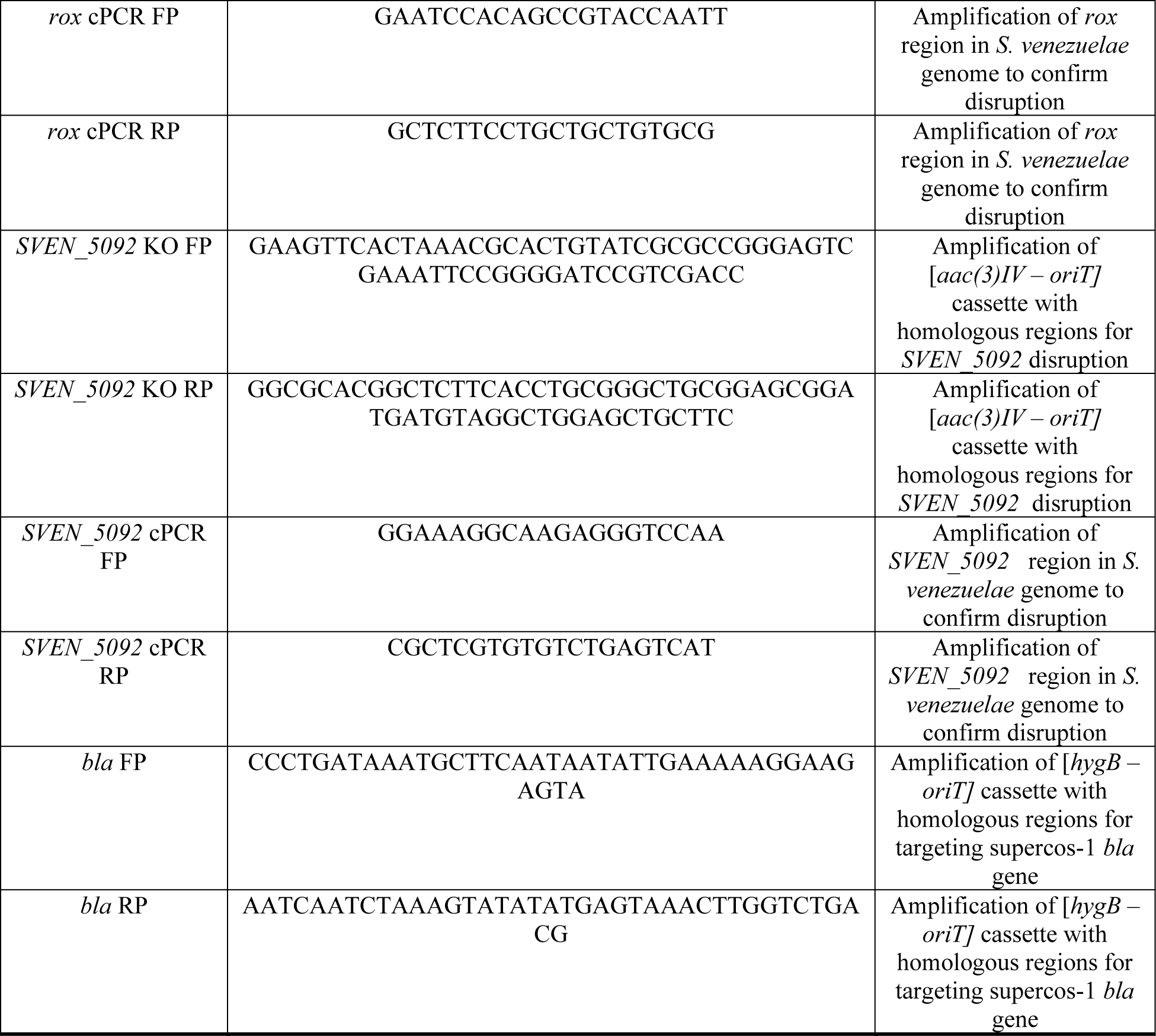

## Oligo list

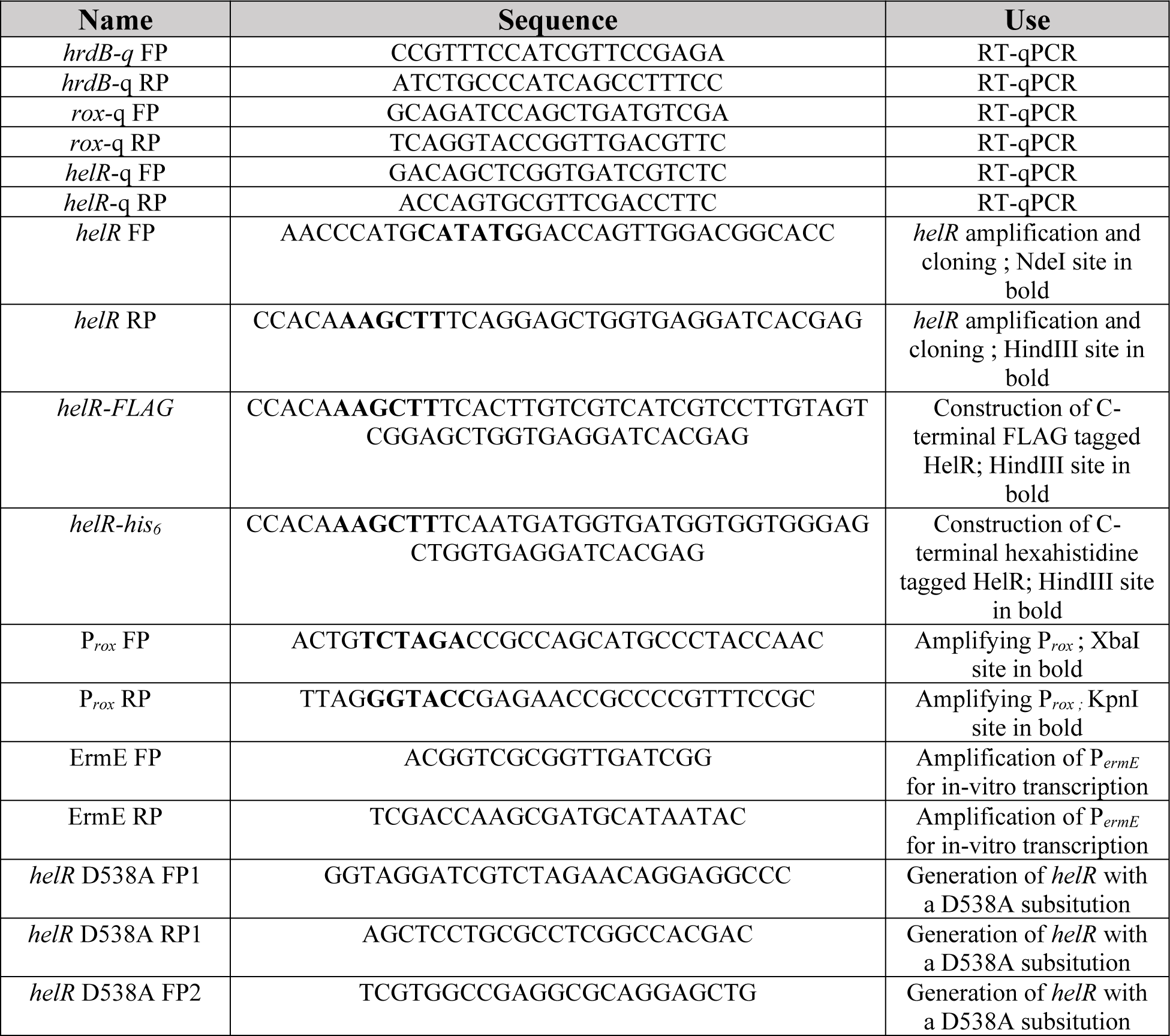
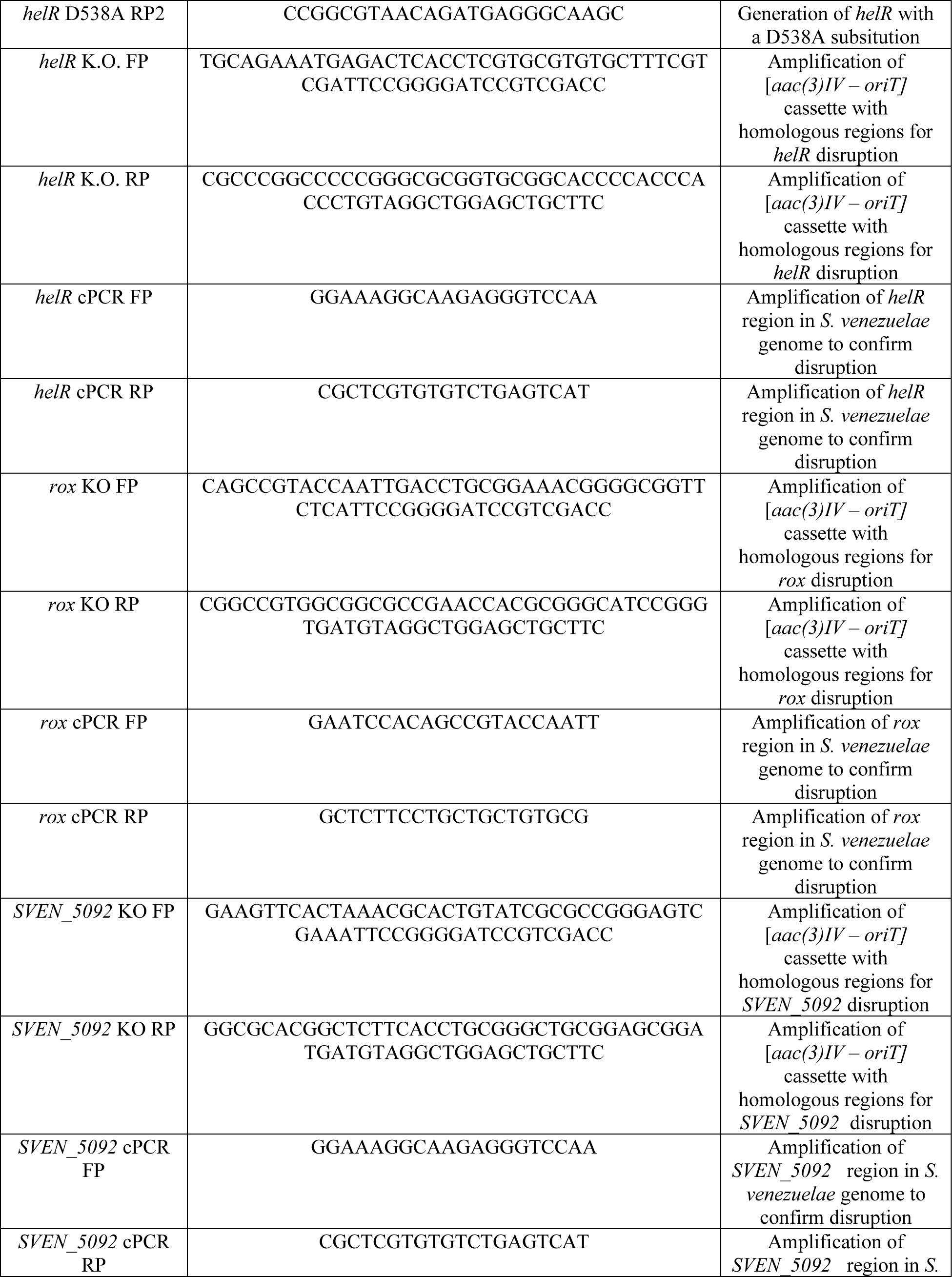
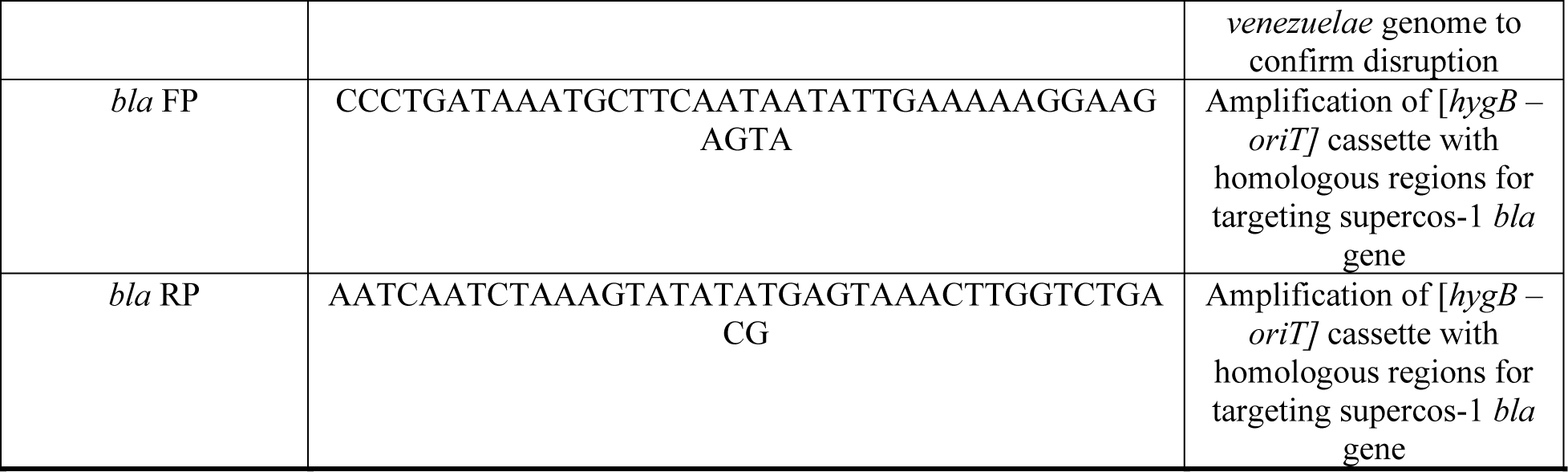

