## Supplemental material for "HelR is a helicase-like protein that protects RNA polymerase from rifamycin antibiotics"

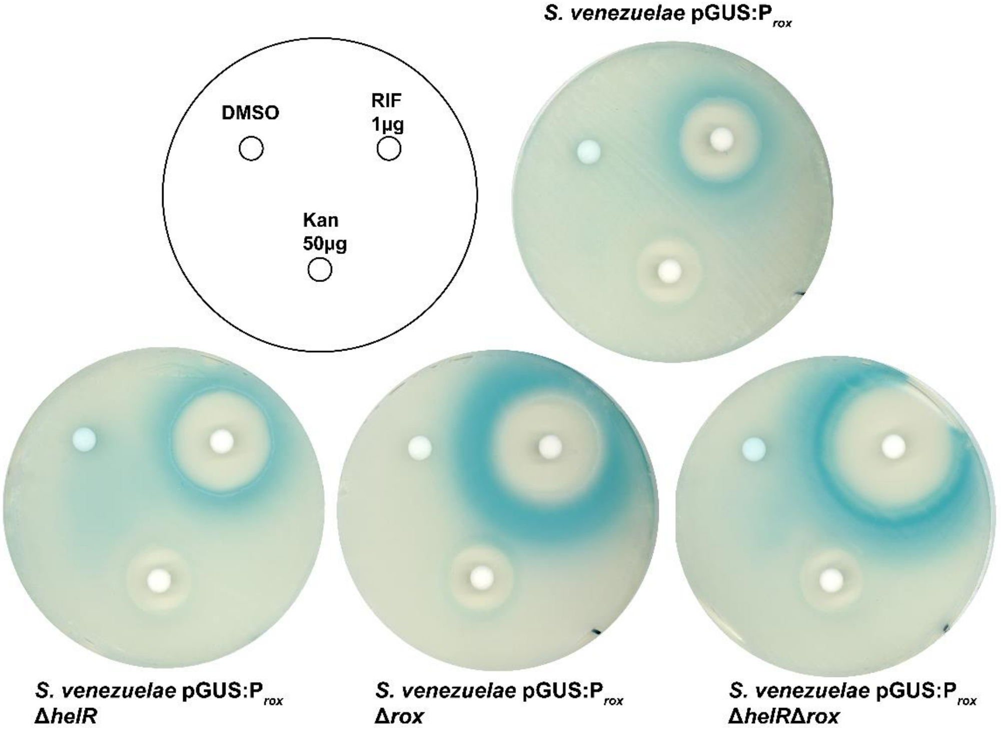


**Supplementary Figure 1 Induction through the RAE does not require HelR** *S. venezuelae* mutants transformed with pGUS:$P_{rox}$ were streaked for confluence on media containing colorimetric GUS substrate. Zones of blue indicate GUS activity and therefore induction of $P_{rox}$. Although some strains are more susceptible than others all are induced by sub-inhibitory rifampin. *S. venezuelae* Δ*rox* and Δ*rox*Δ*helR* appear to have the higher expression in response to rifampin. We hypothesize this is because cells that possess *rox* can inactivate rifampin, dampening induction caused by these compounds over time.

**Table S1 Top 10 Enriched and Depleted RNAP associated proteins**

| Protein | Annotation / Predicted function | Fold Enrichment |
|---|---|---|
| HelR | Superfamily 1 helicase | 426 |
| SVEN_4182 | TetR Family transcriptional regulator | 102 |
| RplQ | Ribosomal protein L17 | 79 |
| SVEN_5967 | Butyryl-CoA dehydrogenase | 40 |
| SVEN_2381 | DUF2344 domain-containing protein | 19 |
| SVEN_3586 | Pyruvate dehydrogenase E1 beta subunit | 17 |
| SVEN_6463 | Outer membrane protein RomA | 15 |
| GltX | Glutamate tRNA ligase | 11 |
| SVEN_5919 | Ferredoxin sulfite reductase | 10 |
| SVEN_1435 | Putative ABC transporter ATP-binding protein | 8 |
| **Protein** | **Annotation / Predicted function** | **Fold Depletion** |
| SVEN_0613 | Secreted protein (Arabinose binding) | 19.2 |
| SVEN_7112 | SAM-dependent methyltransferase | 14.4 |
| SVEN_4686 | Secreted protein (OmpA-like domain) | 8.5 |
| GrpE | Nucleotide exchange factor for DnaK | 5.3 |
| RpoZ | RNA polymerase ω subunit | 5.3 |
| SVEN_1812 | Cytochrome C oxidase subunit 1 | 4.5 |
| AlaS | Alanine tRNA ligase | 4.1 |
| SVEN_1149 | Enoyl ACP reductase | 3.6 |
| SVEN_3043 | Pyruvate formate-lyase | 3.6 |
| SVEN_2206 | PucR family regulator | 3.2 |

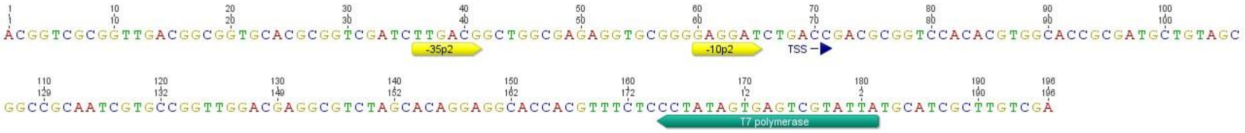


**Supplementary Figure 2 PermE* template** Nucleotide sequence for template DNA used for *in-vitro* transcription in this study. This fragment contains a promoter for T7 RNA polymerase, although this functionality was not utilized here.

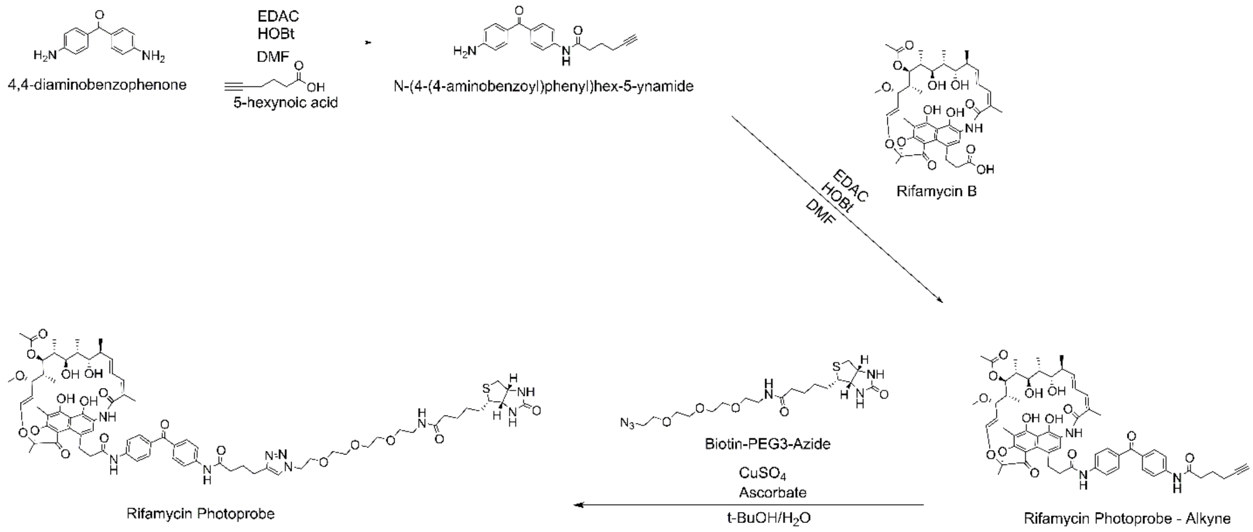


**Supplementary Figure 3 Synthesis of RPP.** Carbodiimide-mediated coupling of 4,4-diaminobenzophenone with 5-hexynoic acid yields an alkyl-terminating aminobenzophenone. Coupling of this reagent to rifamycin B yield a photoactive rifamycin antibiotic. Huisgen cycloaddition (click chemistry) with commercial azide tagged biotin generates the final Rifamycin Photoprobe (RPP).

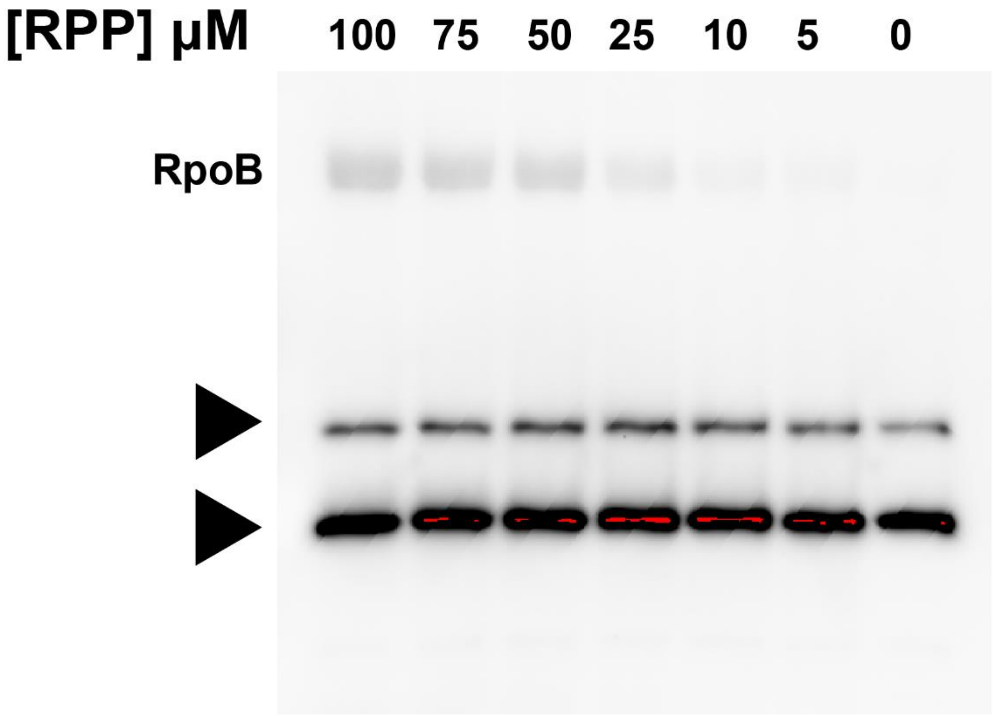


**Supplementary Figure 4 Endogenously biotinylated proteins contaminants** The full blot from Figure 4C. RNAP was incubated with RPP at various concentrations, crosslinked using long wave U.V. light. Protein was separated using SDS-PAGE and analyzed by western blot with Streptavidin-HRP. Black triangles denote endogenously biotinylated proteins in our RNAP preparations. Fortunately these proteins migrated at ~55 and ~70kDa and therefore did not interfere with our detection of labelled RpoB.

**Table S2 Sequence Clusters of BlastP Hits Associated With a RAE**

| Cluster No. | No. Sequences | No. Sequences with RAE | Fraction with RAE | Description |
|---|---|---|---|---|
| 8 | 1 | 1 | 1 | Gtf-HelD fusion |
| 12 | 1 | 1 | 1 | Gtf-HelD fusion |
| 53 | 12 | 10 | 0.83 | HelD |
| 68 | 433 | 330 | 0.76 | HelD |
| 74 | 657 | 285 | 0.43 | HelD |
| 124 | 1 | 1 | 1 | HelD |
| 133 | 30 | 25 | 0.83 | HelD |
| 195 | 566 | 2 | 0 | HelD |
| 227 | 3 | 3 | 1 | HelD_partial |
| 230 | 24 | 1 | 0.04 | HelD |
| 235 | 1459 | 1014 | 0.69 | HelD |
| 239 | 5 | 2 | 0.4 | HelD_partial |
| 243 | 1 | 1 | 1 | HelD_partial |
| 247 | 1 | 1 | 1 | HelD_partial |
| 258 | 1 | 1 | 1 | HelD_partial |
| 259 | 1 | 1 | 1 | HelD_partial |
| 266 | 83 | 63 | 0.76 | HelD |
| 270 | 3 | 3 | 1 | HelD_partial |
| 280 | 1 | 1 | 1 | HelD_partial |
| 313 | 1 | 1 | 1 | HelD_partial |
| 319 | 3 | 2 | 0.67 | HelD_partial |
| 342 | 3 | 1 | 0.33 | HelD_partial |
| 346 | 82 | 36 | 0.44 | Gtf |
| 354 | 7 | 6 | 0.86 | HelD_partial |
| 361 | 237 | 145 | 0.61 | Gtf |
| 367 | 332 | 134 | 0.4 | Gtf |
| 370 | 23 | 7 | 0.3 | Gtf |
| 381 | 7 | 1 | 0.14 | HelD_partial |
| 384 | 5 | 5 | 1 | HelD_partial |
| 391 | 1 | 1 | 1 | HelD_partial |
| 405 | 2 | 2 | 1 | HelD_partial |
| 414 | 1 | 1 | 1 | HelD_partial |
| 417 | 1 | 1 | 1 | HelD_partial |

Gtf = Glycosyltransferase

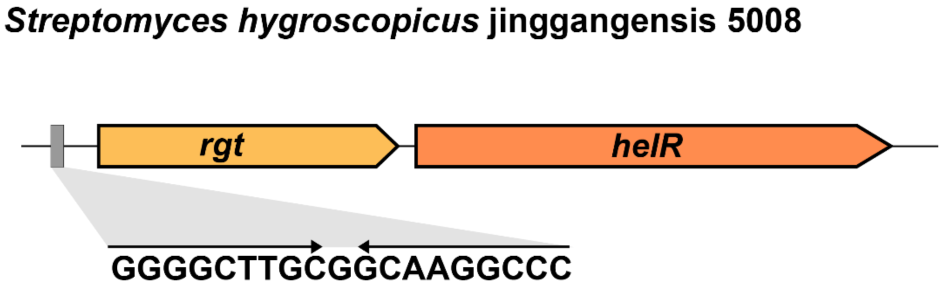


**Supplementary Figure 5 *rgt-helR* operon** The glycosyltransferase sequences associated with a RAE from Table S2 were identified as Rgt, the rifampin glycosyltransferase (Spanogiannopoulos et al. 2012). These were pulled into this dataset due to the presence of predicted glycosyltransferase-HelR fusions. We have previously noted the presence of a single RAE controlling the expression of *rgt* and *helR* in several species, one such example is shown above. Because the number of Rgt-HelR fusions is scant (n = 2) we believe these are errors arising from sequencing/assembly of these operons and are not true fusion proteins. These fusion proteins were unintentionally used to query Refseq genomes which resulted in many hits for Rgt. We therefore removed all the glycosyltransferase protein clusters from our analysis.

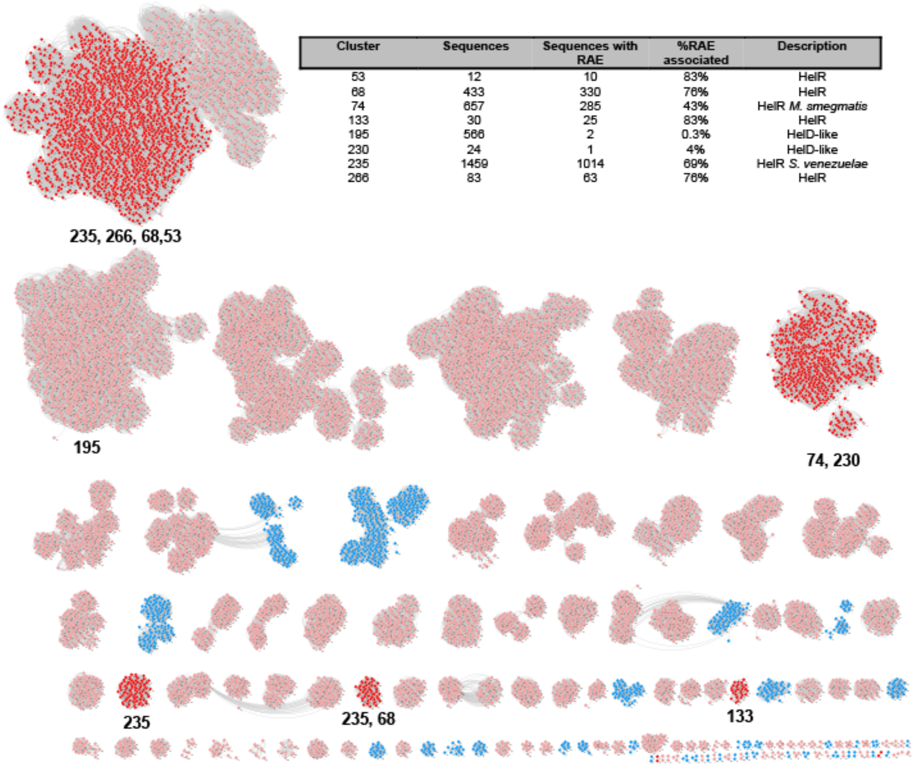


| Cluster | Sequences | Sequences with RAE | %RAE associated | Description |
|---|---|---|---|---|
| 53 | 12 | 10 | 83% | HelR |
| 68 | 433 | 330 | 76% | HelR |
| 74 | 657 | 285 | 43% | HelR *M. smegmatis* |
| 133 | 30 | 25 | 83% | HelR |
| 195 | 566 | 2 | 0.3% | HelD-like |
| 230 | 24 | 1 | 4% | HelD-like |
| 235 | 1459 | 1014 | 69% | HelR *S. venezuelae* |
| 266 | 83 | 63 | 76% | HelR |

**Supplementary Figure 6 Distribution of HelR clusters in the Sequence Similarity Network of HelD-like proteins** The SSN from Figure 11 with the protein cluster identities noted on the network.

**Supplementary Figure 7 Rifamycin Photo Probe (RPP) NMR assignment**

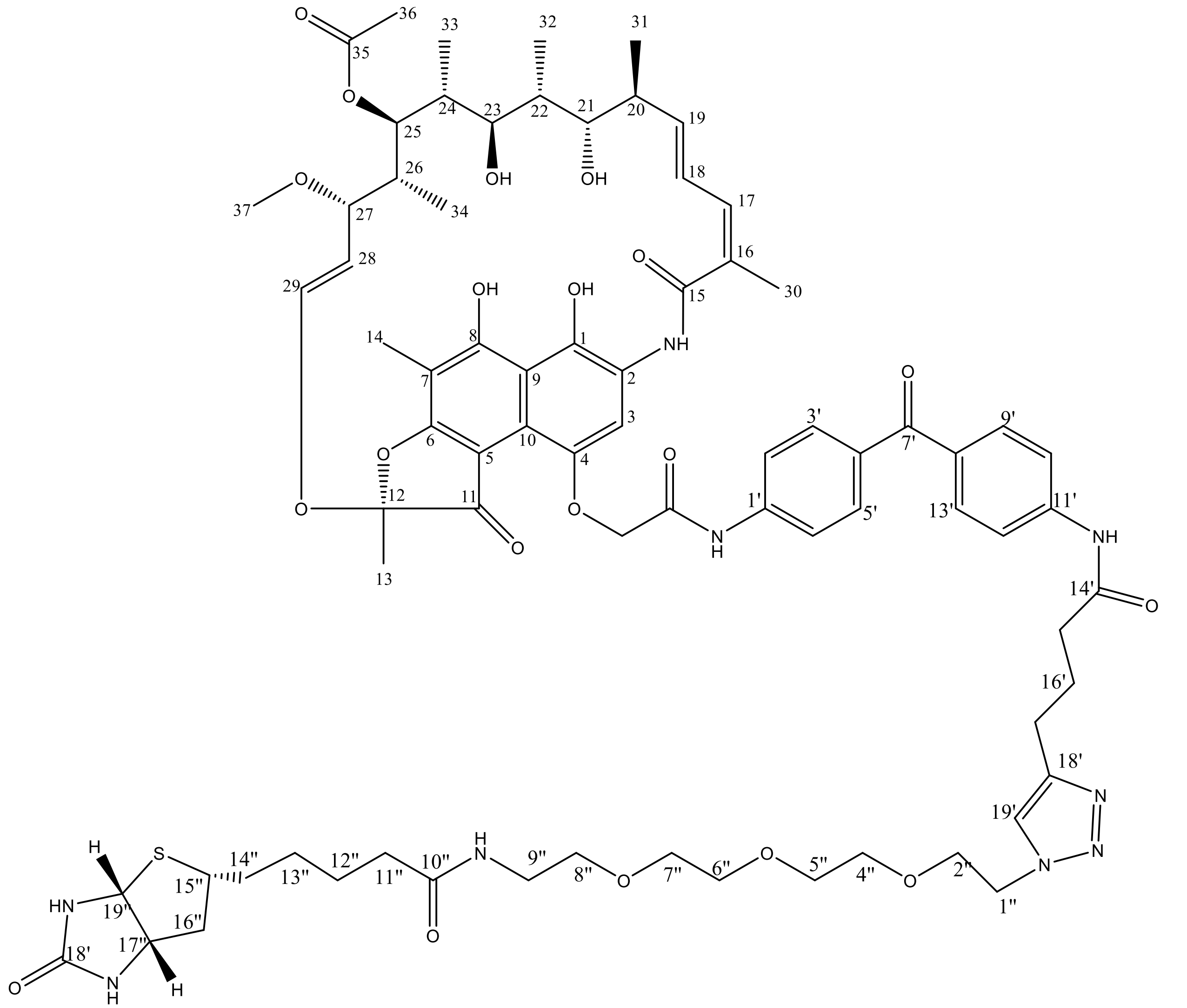

**Table S3:** Chemical shifts for Rifamycin B and Rifampin Photo Probe in dmso-d6, reported in ppm.

| # | $^{1}H$ **Rif B** | $^{13}C$ **Rif B** | $^{1}H$ **RPP** | $^{13}C$ **RPP** |
|---|---|---|---|---|
| 1 | - | 145.17 | | 146.17 |
| 2 | - | 119.37 | - | 119.05 |
| 3 | 7.39 (s, 1H) | 108.71 | 7.67 (s, 1H) | 107.62 |
| 4 | - | *145.34* | - | 142.60 |
| 5 | - | 114.07 | - | 114.51 |
| 6 | - | 171.96 | - | 169.60 |
| 7 | - | 100.26 | - | 99.51 |
| 8 | - | 114.07 | - | 114.51 |
| 9 | - | 115.88 | - | 114.51 |
| 10 | - | 98.72 | - | 99.83 (HMBC) |
| 11 | - | 184.53 | - | 184.37 (HMBC) |
| 12 | - | 108.73 | - | 107.62 (HMBC) |
| 13 | 1.60 (s, 3H) | 22.14 | 1.68 (s, 3H) | 22.36 |
| 14 | 2.01 (s, 3H) | 7.16 | 1.95 (s, 3H) | 7.48 |
| 15 | - | 168.88 | - | 169.70 |
| 16 | - | 130.85 | - | 130.88 |
| 17 | 6.19 (m. 1H) | 131.34 | 6.14 (m, 1H) | 130.81 |
| 18 | 6.35 (d, J = 14.5 Hz, 1H) | 124.6 | 6.44 (m, 1H) | 122.30 |
| 19 | 5.91 (s, 1H) | 139.17 | 5.95 (m, 1H) | 138.45 |
| 20 | 2.14 (m, 1H | 37.58 | 2.15 (m, 1H) | 38.42 |
| 21 | 3.68 (s, 1H) | 71.64 | 3.69 (m, 1H) | 73.23 |
| 22 | 1.56(m, 1H) | 32.25 | 1.59 (m, 1H) | 33.4 |
| 23 | 2.88 – 2.82 (m, 1H) | 75.88 | 2.80 (m, 1H) | 75.95 |
| 24 | 1.22 (m, 1H) | 37.48 | 1.26 (m, 1H) | 37.65 |
| 25 | 4.99 (d, J = 10.7 Hz, 1H) | 72.9 | 5.01(m, 1H) | 73.35 |
| 26 | 0.88 (m, 1H) | 39.47 | 0.88 (m, 1H) | 40.21 |
| 27 | 3.21 (m, 1H) | 76.04 | 3.17 (m, 1H) | 75.41 |
| 28 | 4.89 (s, 1H) | 117.65 | 4.93 (m, 1H) | 117.2 |
| 29 | 6.16 (m, 1H) | 142.52 | 6.18 (m, 1H) | 143.10 |

| | | | | |
|---|---|---|---|---|
| 30 | 1.94 (s, 3H) | 20.70 | 1.97 (m, 3H) | 20.59 |
| 31 | 0.83 (m, 3H) | 17.58 | 0.87 (m, 3H) | 18.17 |
| 32 | 0.82 (m, 3H) | 11.17 | 0.86 (m, 3H) | 12.14 |
| 33 | 0.40 (s, 3H) | 8.05 | 0.32 (m, 3H) | 8.22 |
| 34 | -0.45 | 9.03 | -0.26 (m, 3H) | 8.69 |
| 35 | - | 172.87 | - | 172.08 |
| 36 | 1.92 (s, 3H) | 19.63 | 1.93 (m, 3H) | 20.01 |
| 37 | 2.84 (s, 3H) | 55.71 | 2.88 (m, 3H) | 55.40 |
| **-CH2-** COOH | 4.55 (d, J = 13.1 Hz, 2H) | 66.53 | 4.47 (t, *J* = 5.3 Hz, 2H) | 49.20 |
| -CH2-**CO-** | | | - | 171.53 |
| NH | 9.57 (s, 1H) | - | ***10.2**** *(s, 1H)* | **-** |
| 1’ | - | - | - | 146.18 |
| 2’/6’ | - | - | 8.15 (m, 1H) | 118.99 |
| 3’/5’ | - | - | 7.73 (m, 1H) | 130.91 |
| 4’ | - | - | - | 131.64 |
| 7’ | - | - | - | 193.46 |
| 8’ | - | - | - | 132.29 |
| 9’/13’ | - | - | 7.79 (m, 1H) | 130.67 |
| 10’/12’ | - | - | 7.78 (m, 1H) | 118.17 |
| 11’ | - | - | - | 146.18 |
| 14’ | - | - | - | 171.53 |
| 15’ | | | 2.68 (t, J = 7.6 Hz, 2H) | 24.56 |
| 16’ | | | 1.96 (m, 2H) | 24.82 |
| 17’ | | | 2.44 (t, J = 7.4 Hz, 2H) | 35.89 |
| 18’ | | | - | nd |
| 19’ | - | - | *7.67 (m, 1H)* | *107.12* |
| 11’-**NH-** | | | 10.28 (s, 1H) | - |
| 1’’ | - | - | 4.47 (t, J = 5.3 Hz, 2H) | 49.25 |
| 2’’ | - | - | 3.80 (t, J = 5.3 Hz, 2H) | 68.76 |
| 3’’ | - | - | 4.28 (m, 2H) | 59.17 |
| 4’’ | - | - | 4.11 (m, 2H) | 61.01 |
| 5’’ | - | - | 3.48 (m, 2H) | 69.56 |
| 6’’ | - | - | 3.52 (m, 2H) | 69.68 |
| 7’’ | - | - | 3.47 (m, 2H) | 69.62 |
| 8’’ | - | - | 3.38 (t, J = 6.0 Hz, 2H) | 69.14 |

| | | | | |
|---|---|---|---|---|
| 9’’ | - | - | 3.17 (q, *J* = 5.8 Hz, 2H) | 38.42 |
| 9’’-**NH**-CO- | | | 7.8 (s, 1H) | - |
| 10’’ | - | - | - | 172.2 |
| 11’’ | - | - | 2.05 (t, J = 7.5 Hz, 2H) | 35.08 |
| 12’’ | - | - | 1.48 (m, 2H) | 25.27 |
| 13’’ | - | - | 1.28 (m, 2H) | 28.02 |
| 14’’ | - | - | 1.43/1.59 (m, 2H) | 28.17 |
| 15’’ | | | 3.08 (m, 2H) | 55.39 |
| 16’’ | - | - | 4.12 (m, 2H) | 61.16 |
| 17’’ | - | - | 4.29 (dd, J = 7.7, 5.2 Hz, 1H) | 59.25 |
| 17’’-N**H**-19’’ | | | 6.34 (s, 1H) | - |
| 18’’ | - | - | - | 162.99 |
| 18’’-N**H**-19’’ | - | - | 6.40 (t, J = 1.9 Hz, 1H) | - |
| 19’’ | - | - | 4.11 (ddd, J = 7.8, 4.5, 2.0 Hz, 1H) | 60.93 |

### NMR Spectra

**Supplementary Figure 8 $^{1}$H NMR spectra of RPP in dmso-$d_6$**

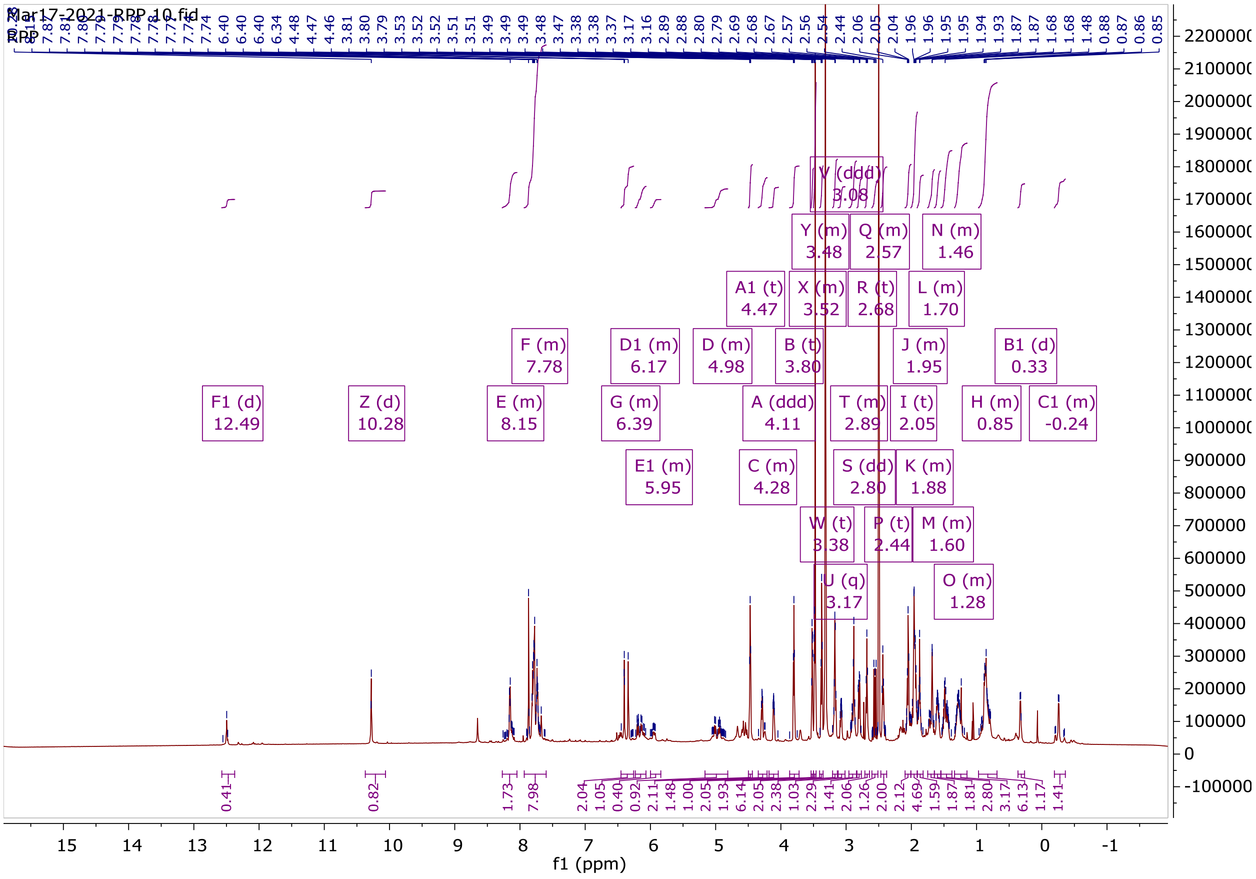

**Supplementary Figure 9 $^{1}H$ NMR spectra overlay of Rifamycin B and RPP in dmso-$d_6$**

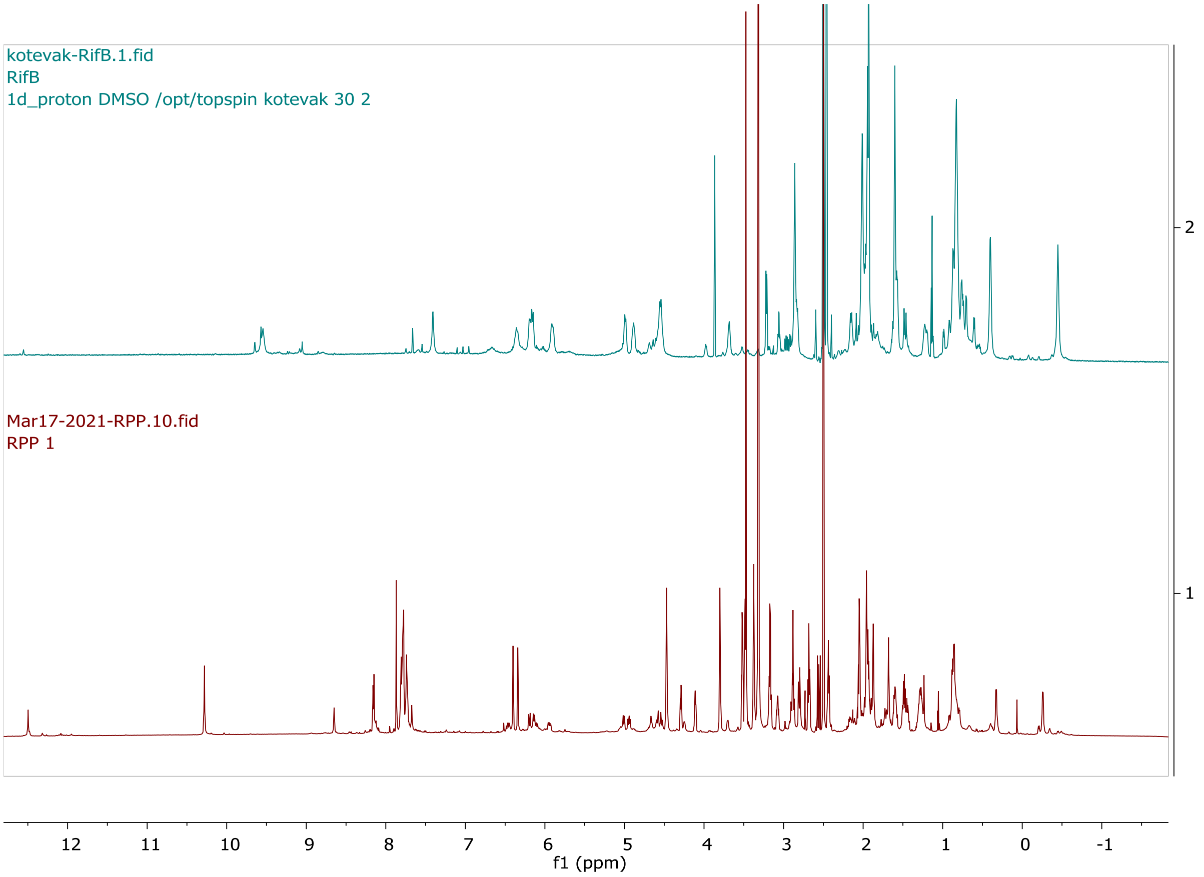

**Supplementary Figure 10 $^{13}C$ NMR spectra of RPP in dmso-$d_6$**

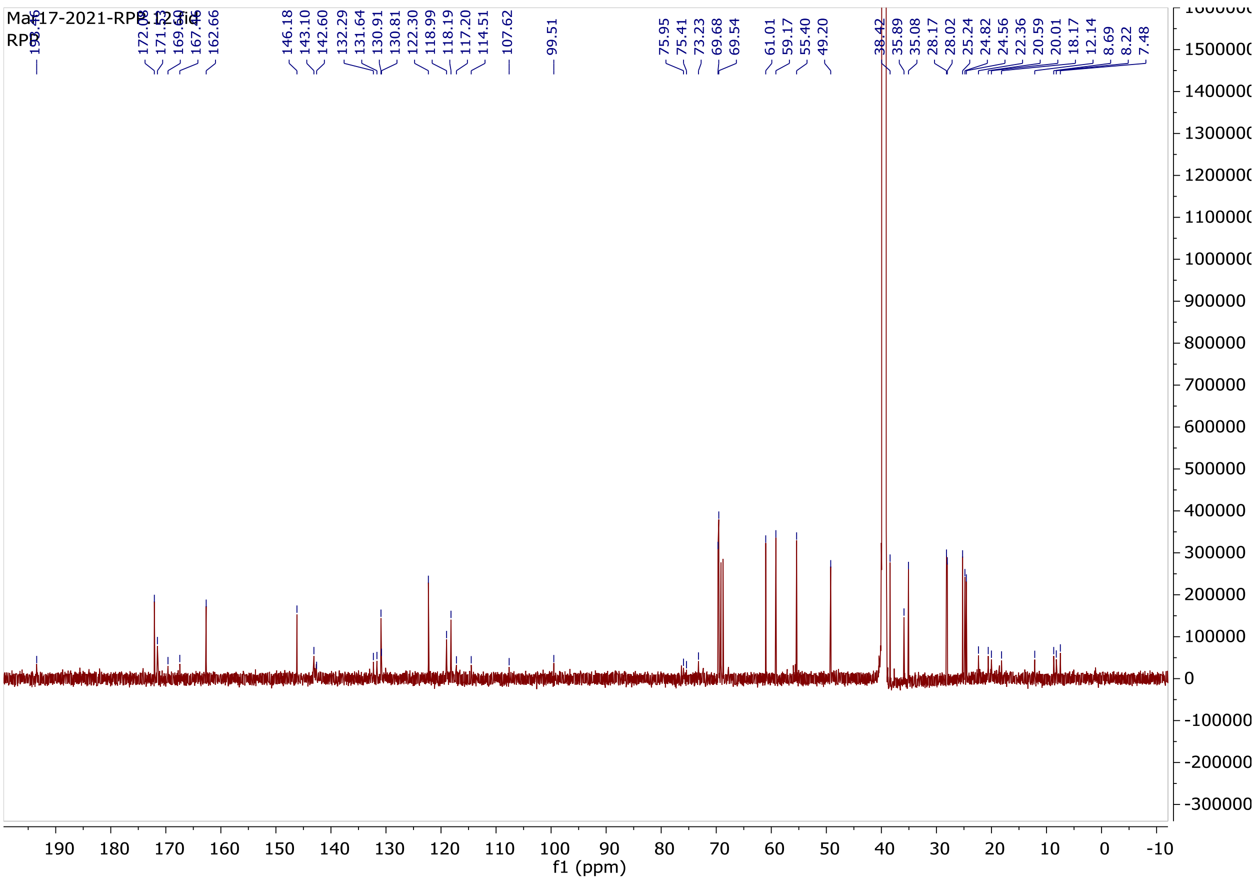

**Supplementary Figure 11 $^{1}H$-$^{1}H$- COSY spectra of RPP in dmso-$d_6$**

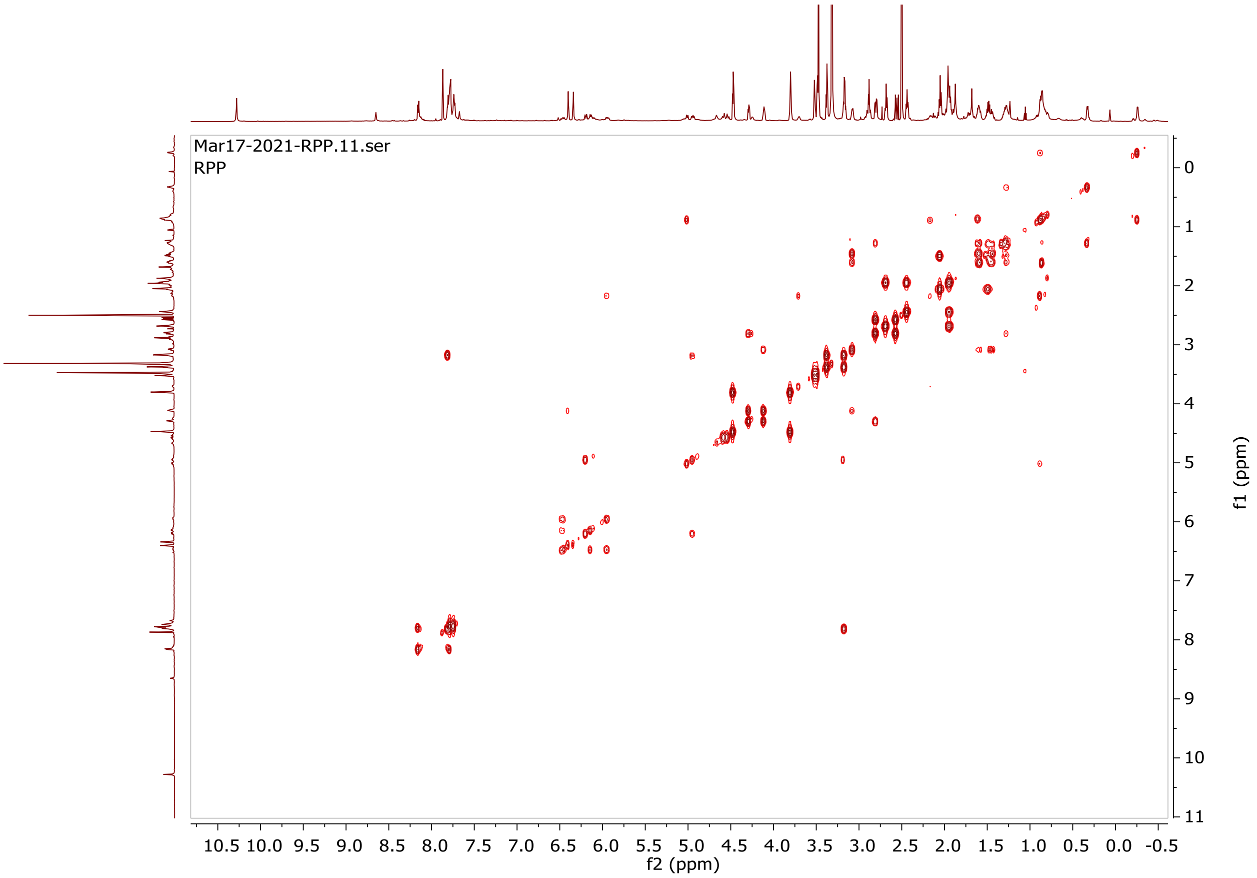

**Supplementary Figure 12 $^{1}H$-$^{13}C$- HSQC spectra of RPP in dmso-$d_6$**

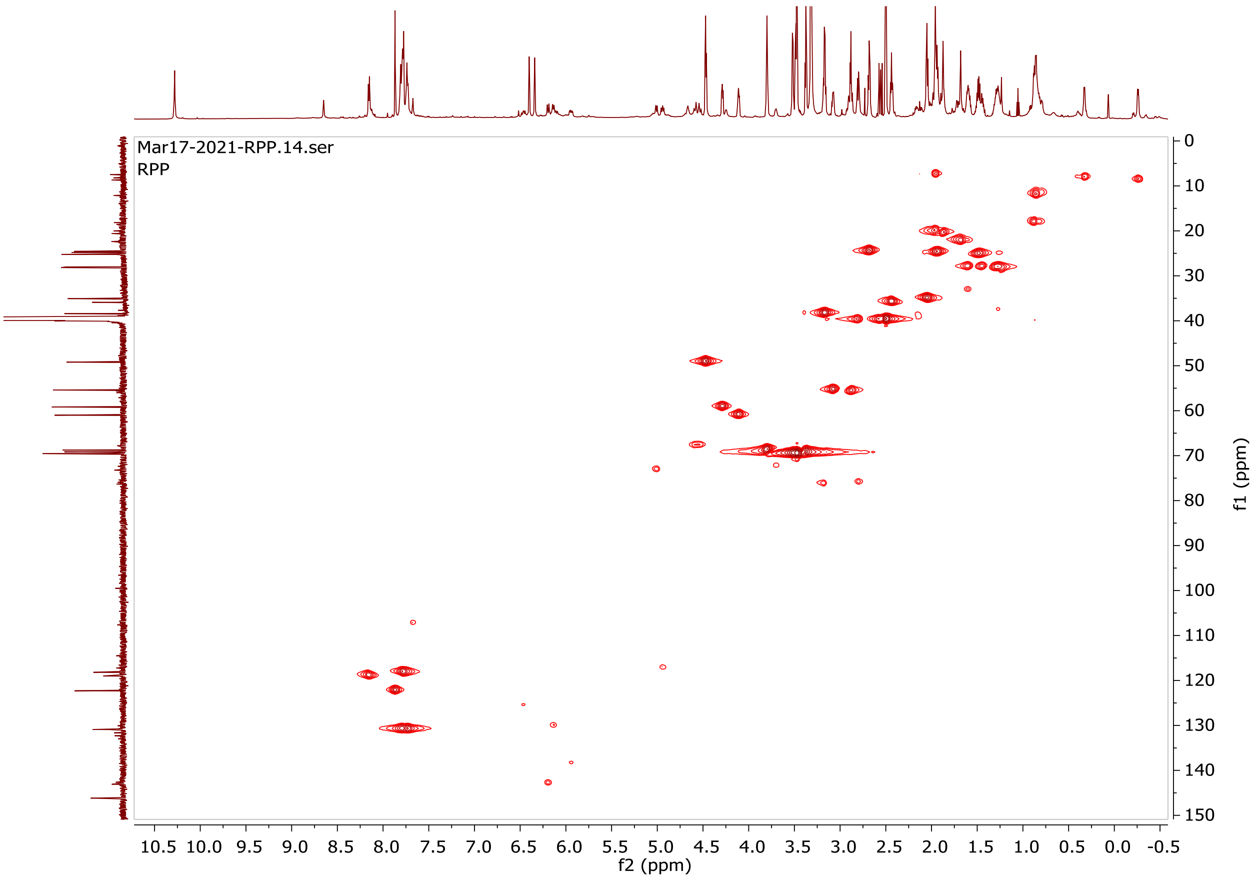

**Supplementary Figure 13 $^{1}H$-$^{13}C$- HMBC spectra of RPP in dmso-$d_6$**

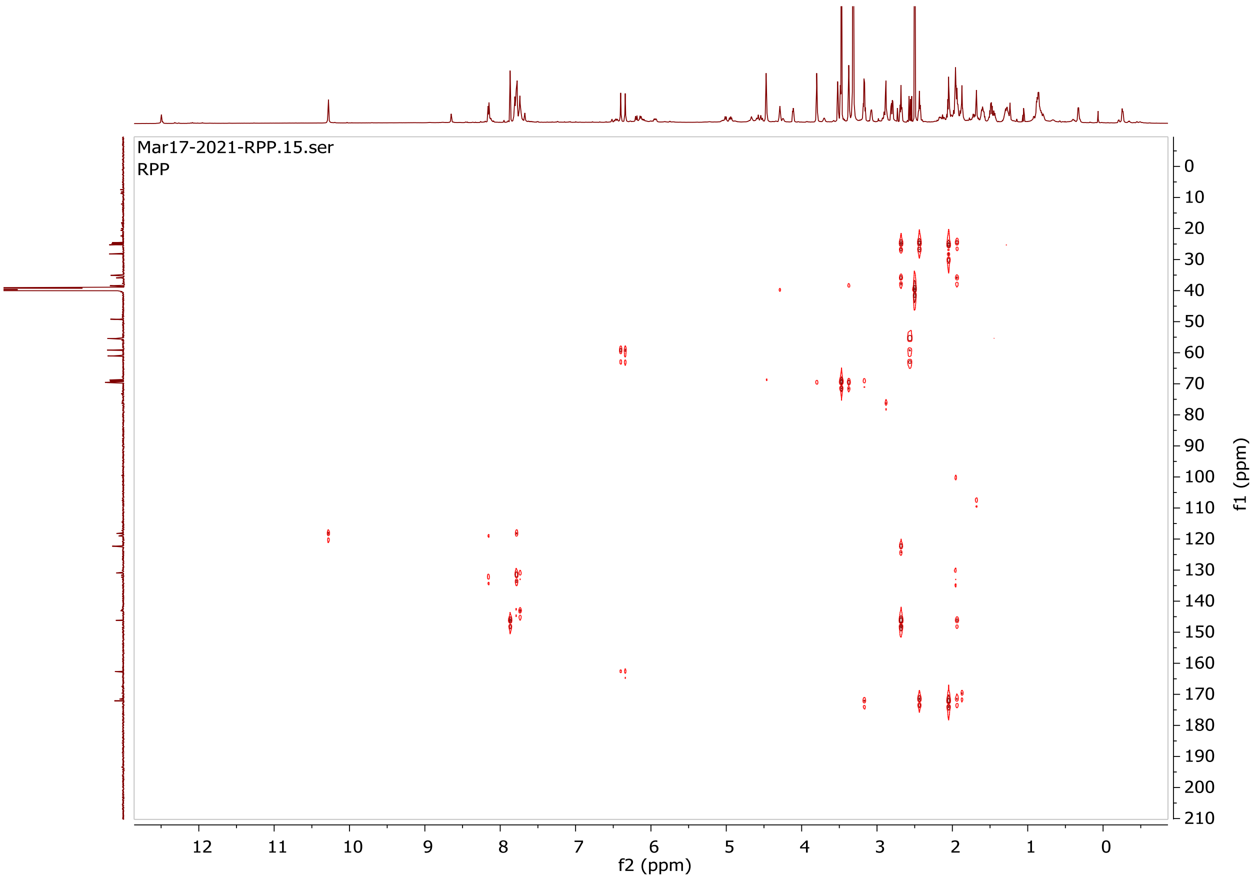

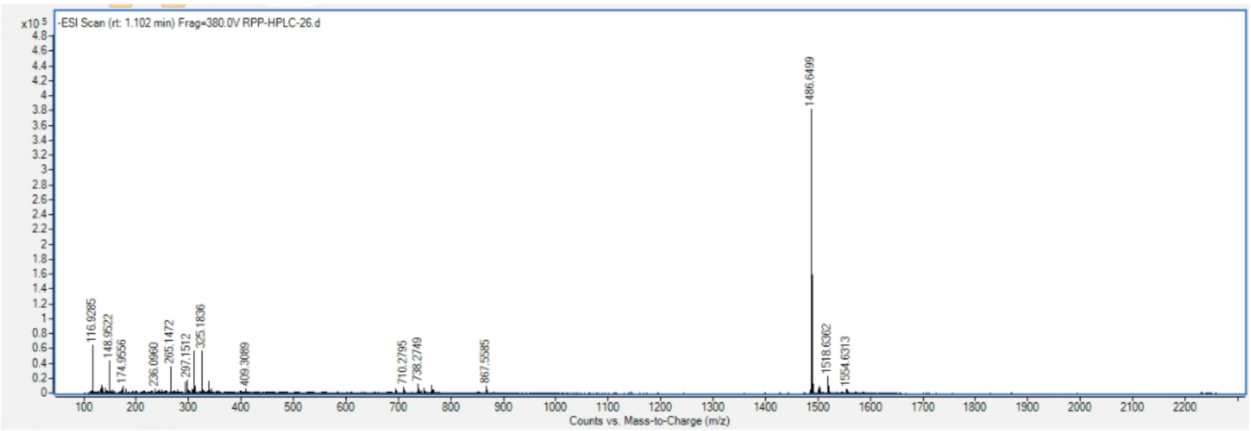


**Supplementary Figure 14 HRMS spectra for RPP** HRMS: Calculated for $C_{76}H_{97}N_9O_{20}S$ 1487.6571 Da, $[M-H]^-$ $C_{76}H_{96}N_9O_{20}S$: 1486.6498 Da, found 1486.6499 Da.
